# Directed Evolution in Codon Space

**DOI:** 10.64898/2026.08.03.742557

**Authors:** James Heuschkel, Laura Kingsley, Jon Reed, Di Li, Matthew Warner, Noah Pefaur, Steven Cramer

## Abstract

Directed evolution is commonly used in protein engineering, where mature molecules are routinely improved through iterative local search of amino acid space. Here, we extend this principle to coding DNA. We developed a language-model-guided framework that iteratively refined industry-optimized coding sequences of clinical-stage therapeutics through synonymous exploration of codon space. Across 23 antibody-based therapeutics, SynCodonLM-guided refinement significantly increased recombinant expression in CHO cells for 17 molecules (74% responder rate), without significant compromise of product-quality or biophysical attributes. Moreover, changes in model likelihood predicted expression gains more effectively than heuristic statistical or mRNA-structure descriptors, despite no explicit expression objective. Codon-level likelihood also tracked temporal progression in influenza A H1N1 sequences, indicating the model captures evolutionary signal. These results show that even production-optimized sequences retain accessible fitness in synonymous codon space, establishing directed evolution as a practical strategy to improve biologic expression, a key manufacturing bottleneck, without altering protein sequence.

## Introduction

The degeneracy of the genetic code creates a large design space in which many synonymous sequences encode the same protein, yet these choices are not functionally equivalent^1,2^. Synonymous codon usage influences translation kinetics^3^, mRNA stability^4^, co-translational folding^5^, protein conformation and function^6^, and product quality^7^ - with downstream consequences in vaccines^8–10^, oncolytic viruses^8^, AAV payloads and capsids^9–11^, and therapeutic antibody yield^12–14^. Synonymous substitutions are therefore not biologically silent.

This complexity has made codon optimization a routine component of recombinant protein and nucleic acid design, including applications in vaccines, gene therapies and therapeutic biologics^15^. Yet codon optimization remains methodologically unsettled. Classical approaches have relied heavily on host-frequency heuristics and related summary statistics, whereas newer machine-learning models generate host-specific coding sequences from protein input using context-aware sequence models^16–22^.

A central limitation is that codon optimization is typically framed as a one-shot design problem, producing a fixed construct. This contrasts with modern protein engineering, where functional molecules are improved through iterative rounds of localized change, and where protein language models have shown that a small number of evolutionarily plausible edits can improve even highly mature antibodies^23^.

Recent advances in codon language modeling suggest that an analogous framework may now be possible directly in coding DNA space^20,24–28^. Language models trained on coding DNA sequences rather than amino acid sequences have been shown to capture biologically informative signals in downstream tasks, indicating that coding sequences contain useful information beyond the protein sequence they encode^26^. Codon language models trained under synonym-constrained prediction further raise the possibility that codon-level sequence modeling can isolate DNA-level constraints from amino-acid-level semantics and thereby provide a principled basis for post-design synonymous refinement^24^.

Here, we show that codon-level likelihood reflects evolutionary structure, significantly tracking temporal progression in several influenza A (H1N1) proteins, establishing that the model encodes a biologically meaningful signal rather than a spurious statistical artifact. Leveraging this encoding, we apply our directed evolution approach to 23 patented, clinically deployed antibody-based therapeutics, treating each disclosed coding sequence as a high-fitness parental baseline for iterative synonymous refinement rather than as a fully optimized endpoint. At each step, a context-aware codon model proposes the synonymous substitution predicted to most improve local sequence compatibility, the sequence is updated, and the modified context is rescored before the next edit is selected. Most importantly, we demonstrate that these clinically validated therapeutic coding sequences remain locally improvable in codon space.

## Results

### Reframing codon optimization as an iterative search process

Whereas protein engineering routinely improves functional molecules through iterative rounds of localized change (**Fig. 1a**), codon optimization is typically performed as a one-time design step (**Fig. 1b**). We therefore reframed synonymous sequence design as an iterative search problem, treating clinically validated coding sequences as high-fitness starting points for further refinement in codon space (**Fig. 1c**).

**Fig. 1:**
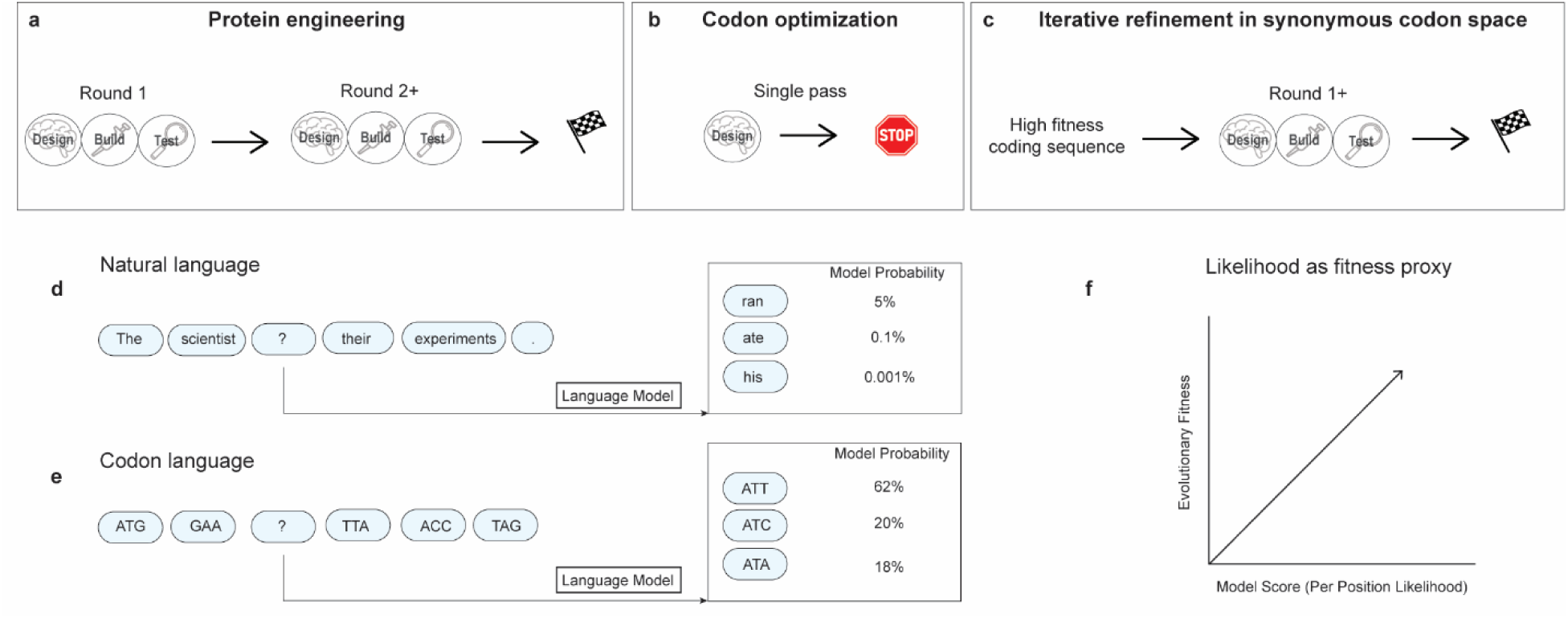
Reframing codon optimization as an iterative, language-model-guided refinement in codon space. (a) Protein engineering improves functional molecules through iterative rounds of design-build-test cycles, progressively enhancing sequence performance. (b) Conventional codon optimization, by contrast, is performed as a single-pass design step, producing a fixed coding sequence that is not revisited. (c) Here, clinically validated coding sequences are treated as high-fitness starting points and iteratively refined in codon space through context-aware synonymous substitutions guided by a codon language model, without altering the encoded protein. (d) A natural-language model assigns probabilities to candidate words conditioned on their surrounding context. (e) Analogously, a codon language model assigns probabilities to synonymous codons conditioned on their local sequence context, scoring a masked position over its synonymous alternatives. (f) Model-assigned likelihood is used as a proxy for evolutionary fitness, consistent with prior work^23,29,30^ showing that language-model likelihoods correlate with sequence fitness in protein engineering.

To guide this search, we drew on an analogy to natural language modeling. Just as a natural-language model assigns probabilities to candidate words based on their surrounding context (**Fig. 1d**), a codon language model assigns probabilities to synonymous codons conditioned on local sequence context (**Fig. 1e**). Because language-model likelihoods have been shown to correlate with sequence fitness in protein engineering^23,29,30^, we used codon-level likelihood as a proxy for sequence compatibility, defining an evolutionary score ("Evo score") for each coding sequence (**Fig. 1f**).

### A codon language model captures evolutionary context

To first validate that SynCodonLM encodes biologically meaningful evolutionary information, we evaluated model-derived Evo scores on an independent viral evolution benchmark spanning Influenza A (H1N1) nucleoprotein, hemagglutinin, and neuraminidase coding sequences. Critically, this analysis was performed without task-specific fine-tuning, enabling direct assessment of whether codon-level likelihood alone reflects evolutionary trajectories.

Across all three proteins, SynCodonLM Evo scores increased significantly with sampling time (r = 0.41–0.79; all *p* < 1e-3; **Fig. 2a, c, e**), indicating that sequences observed later in viral evolution are assigned higher model compatibility. Nucleoprotein additionally showed a clear temporal shift, coincident with the 2009 pandemic strain A/California/07/2009, followed by sustained elevation in subsequent years. We also evaluated full-vocabulary (non-synonym-constrained) Evo scores (**Supplementary Fig. 1–3**). The temporal signal was reproduced for nucleoprotein and hemagglutinin, but was absent for neuraminidase, which retained a significant positive trend only under synonym-constrained scoring. This indicates that the synonym constraint can be necessary to isolate the codon-level evolutionary signal from amino-acid-level variation.

**Fig. 2:**
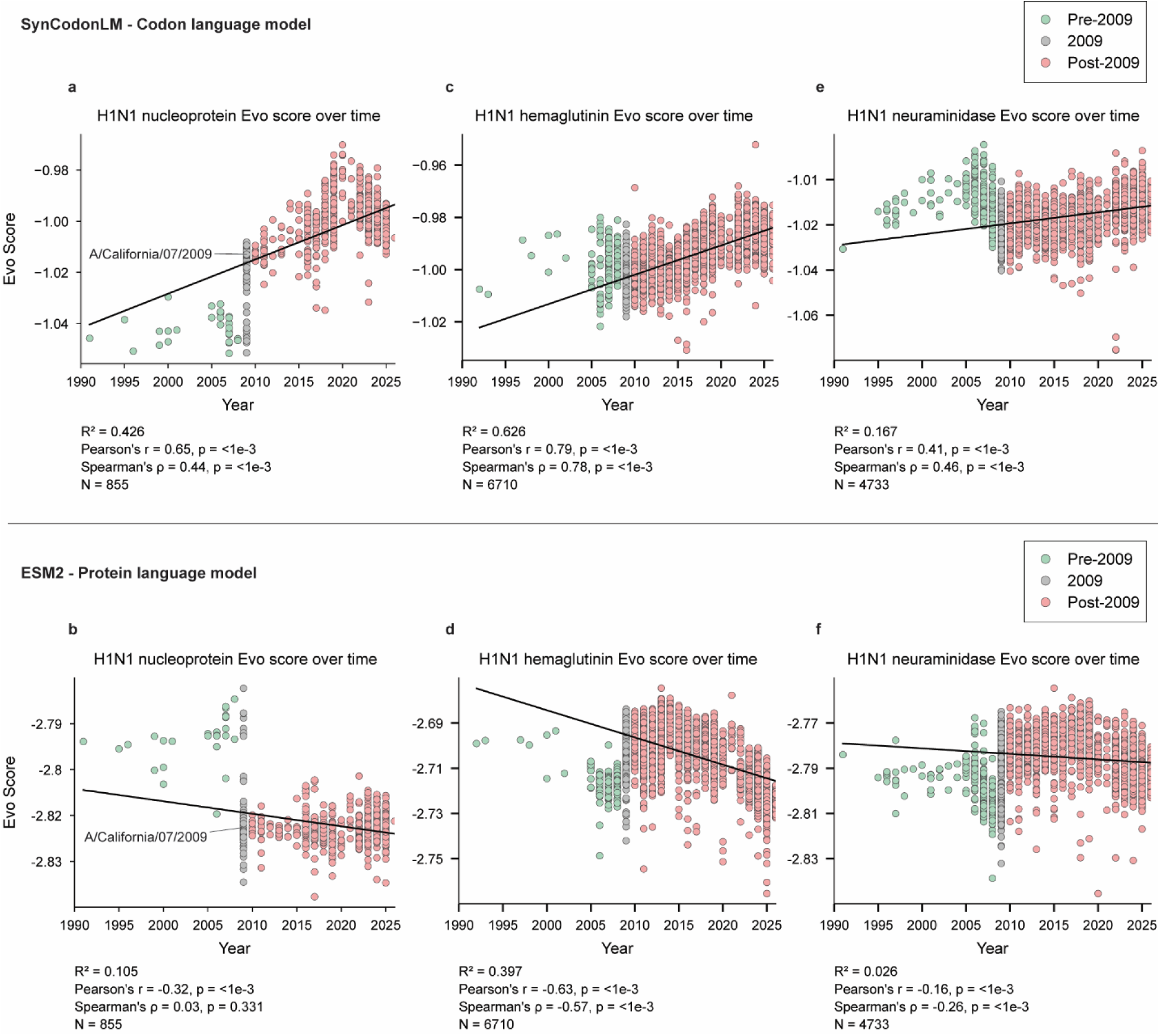
Codon-level likelihood tracks forward viral evolution, whereas protein-level likelihood trends in the opposite direction. (a, c, e) SynCodonLM Evo scores plotted against sampling year for Influenza A H1N1 nucleoprotein (a), hemagglutinin (c), and neuraminidase (e) coding sequences. Scores increase significantly with time in all three proteins (Pearson r = 0.65, 0.79, and 0.41, respectively; all *p* < 1 × 10^−3^), indicating that later-emerging sequences are assigned higher model likelihood. Nucleoprotein shows a pronounced shift, coincident with the 2009 pandemic strain A/California/07/2009, sustained in subsequent years. (b, d, f) ESM-2 pseudo-log-likelihoods on the matched translated sequences show the opposite trend (r = −0.32, −0.63, and −0.16; all *p* < 1 × 10^−3^), decreasing over time. All sequences correspond to unique viral isolates with non-synonymous diversity; purely synonymous variants were excluded. No task-specific fine-tuning was applied to either model.

**Fig. 3:**
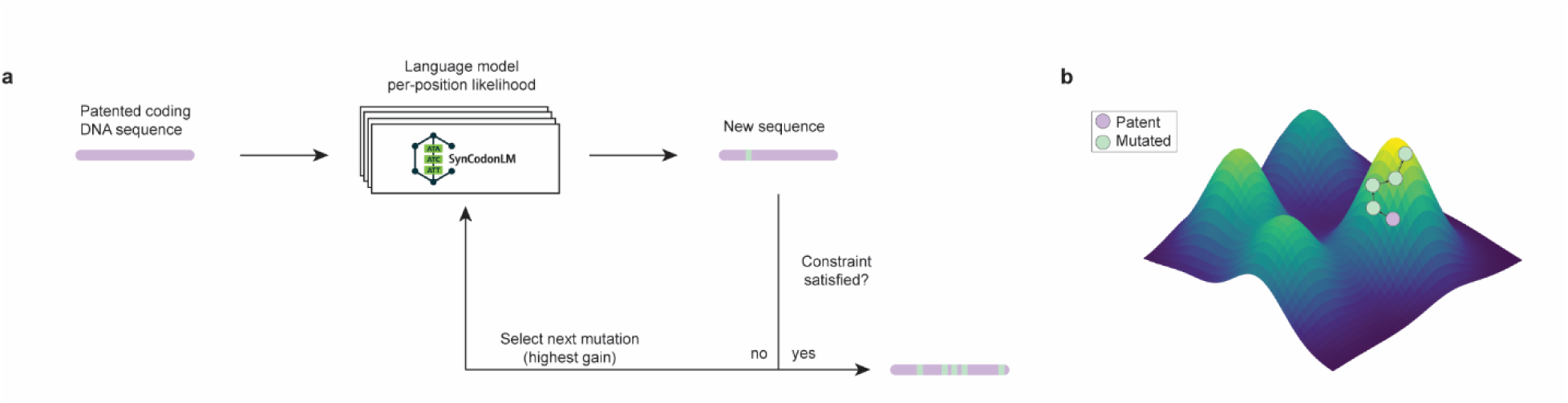
A language-model-guided framework for directed evolution in codon space. (a) Patented antibody coding sequences are passed to SynCodonLM^24^, and the substitution predicted to most increase local likelihood is applied at each iteration. The newly generated sequence is passed back into the model for finding the next most preferable mutation, up to a set limit of mutations or when the sequence is at a maxima. (b) Directed evolution drives the sequence up a local fitness gradient in codon space, defined as Evo score, while preserving the encoded protein. Directed evolution proceeded by repeatedly selecting the substitution predicted to most improve local sequence compatibility, updating the context, and re-scoring before the next edit (Fig. 3a). This allowed successive substitutions to interact through changing context, traversing codon space toward higher-fitness configurations without altering the encoded protein (Fig. 3b).

In contrast, ESM-2 exhibited the opposite temporal behavior on the matched translated sequences (r = −0.16 to −0.63; all *p* < 1e-3; **Fig. 2b, d, f**).

While absolute score scales are not directly comparable due to differences in tokenization and training objectives, the directionality of these trends is striking; codon-level modeling captures a forward-in-time signal that is not recoverable from the matched protein sequences, where ESM-2 likelihood instead trends opposite to sampling year.

Importantly, all sequences analyzed corresponded to distinct viral isolates with non-synonymous diversity, with purely synonymous variants excluded to prevent artificial inflation of SynCodonLM performance relative to protein language models. Despite this conservative design, SynCodonLM systematically assigned higher likelihood to sequences that emerged later in viral evolution.

These findings demonstrate that a simple, unsupervised codon-level likelihood score contains substantial predictive information about real-world viral evolution, indicating that synonymous codon usage encodes context-dependent biological constraints learnable by a language model. If these learned constraints reflect genuine functional information rather than an artifact of natural sequence statistics, then optimizing sequences toward higher codon-level likelihood should yield measurable biological benefits. Therefore, we next ask whether SynCodonLM-guided synonymous edits improve recombinant protein expression.

### A language-model-guided framework for directed evolution in codon space

We developed a language-model-guided framework for directed evolution in synonymous codon space, treating clinically validated coding sequences as high-fitness parental baselines and iteratively refining them while preserving the encoded protein.

To test this framework, we assembled a panel of 23 patented, clinically deployed antibody-based therapeutics and treated their disclosed coding sequences as high-fitness parental baselines. The panel was deliberately constructed to span industrially relevant format and isotype diversity, including conventional monospecific IgGs, bispecific architectures, an antibody–drug conjugate, antibody fusion proteins, and molecules with significant Fc engineering (**Supplementary Table 1**). SynCodonLM-guided refinement was capped at 10% of codon positions per sequence, limiting divergence from the clinically validated parent while allowing sufficient synonymous change to test whether local codon-space refinement could produce measurable effects.

### SynCodonLM-refined sequences increase recombinant antibody titers

We next tested whether iterative synonymous refinement produced measurable changes in recombinant antibody expression. Parental (patented) and SynCodonLM-refined coding sequences were cloned into pTT5-based mammalian expression vectors^31^ with a fixed signal peptide and stop codon and transiently expressed in CHO-3E7^32^ cultures in biological triplicate. After six days, culture supernatants were harvested for titer quantification, and expressed antibodies were purified for downstream analytical characterization.

Importantly, across the panel of 23 clinical-stage antibody-based therapeutics, SynCodonLM-refined sequences increased secreted antibody titer relative to their parental patented counterparts in 17 of 23 cases (74% responder rate, 15 individually significant, paired t-test *p* < 0.05), corresponding to a mean (geometric) 1.34-fold increase (**Fig. 4a**; paired t-test *p* < 0.001; Wilcoxon signed-rank *p* < 0.001). The magnitude of positive responses substantially exceeded that of the few negative ones (**Fig. 4a**). Raw titers are provided in **Supplementary Fig. 4**.

**Fig. 4:**
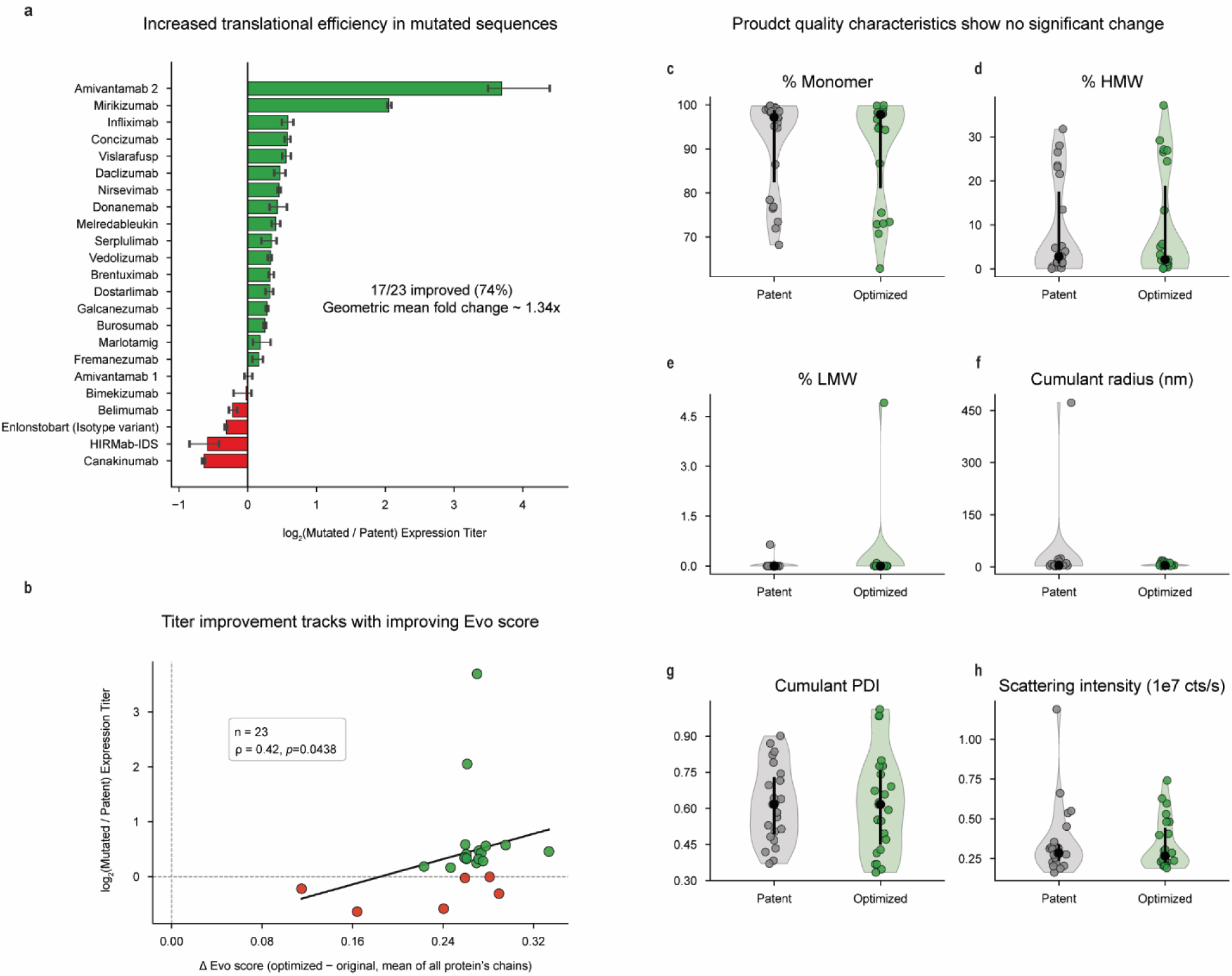
Clinically optimized therapeutic coding sequences remain improvable without compromising product quality. (a) Per-molecule log₂-fold change in secreted CHO titer for SynCodonLM-refined sequences relative to patented parental counterparts across 23 clinical-stage antibody-based therapeutics (mean ± SD, biological triplicates). Green, improved constructs (17/23; 74% responder rate, 15 individually significant by paired t-test, *p* < 0.05); red, reduced-titer constructs. Panel-wide mean (geometric) fold increase ∼1.34 (paired t-test *p* < 0.001; Wilcoxon signed-rank *p* < 0.001). (b) Δlog₂(titer) versus Δ Evo score (refined − parental, averaged across expressed chains); Spearman’s rank correlation coefficient (ρ) = 0.42, *p* = 0.0438, n = 23. All 23 constructs show positive Δ Evo score, confirming that refinement systematically raised model likelihood. (c–e) Analytical size-exclusion chromatography (aSEC) of Protein A–purified material: percent monomer (c), high-molecular-weight species (%HMW, d), and low-molecular-weight species (%LMW, e), for parental (gray) and refined (green) groups; violin plots with median lines. Distributions show no significant difference between groups (paired Wilcoxon signed-rank, *p* > 0.05 for all measures). (f–h) Dynamic light scattering (DLS) of the same material: cumulant hydrodynamic radius (f), polydispersity index (g), and scattering intensity (h). Again, distributions are closely matched between groups, indicating no detectable change in aggregation propensity or homogeneity (paired Wilcoxon signed-rank, *p* > 0.05).

To ask whether these titer changes tracked with the model’s own compatibility signal, we compared the change in mean chain-level Evo score (ΔEvo = refined − parental, averaged across all expressed chains of a given molecule) with the observed log₂-fold change in titer (**Fig. 4b**). Across the panel, Δ Evo score was significantly positively correlated with titer improvement (ρ = 0.42, *p* = 0.0438, n = 23), and all 23 constructs, including the negative responders, showed positive ΔEvo values, confirming that iterative refinement systematically raised codon-language-model likelihood as intended. This relationship indicates that the magnitude of model-implied improvement provides genuine, quantitative information about the likelihood of a titer gain. Chain-resolved correlations were stronger for heavy chains (**Supplementary Fig. 5**), suggesting that heavy-chain optimization may contribute disproportionately to expression gains under the transient expression conditions used here.

Because synonymous changes can affect co-translational folding^5^ and post-translational modifications^7^, we confirmed product identity by intact LC-MS on Protein A–purified material. Each pair of molecules (parental and refined) was evaluated by LC-MS at both the intact and subunit levels to (1) verify sequences and complexation, and (2) gauge the impact(s) on the distribution of abundant PTMs (ex., glycosylation, C-terminal lysine truncation) that comprise the majority of observed proteoforms.

For all molecule pairs, intact and subunit molecular weights matched one another at the global level (per molecule conditions, **Supplementary Table 2**; spectra and summary, **Supplementary Data)**, ruling out coincidental modifications or subtle sequence deviations as sources of the observed titer differences. Additionally, PTM distributions were nearly identical within each pair, but dissimilar across the different molecules, indicating that the amino acid sequence (and not codon differences within the nucleotide sequences) is the major driver for proteoform distribution.

Consistent with this preserved molecular identity, biophysical characterization of the purified material showed no significant adverse shifts in product-quality attributes. Analytical size-exclusion chromatography (aSEC) revealed comparable distributions of monomer content (**Fig. 4c**), high-molecular-weight species (**Fig. 4d**), and low-molecular-weight species (**Fig. 4e**) between parental and refined groups. Complementary dynamic light scattering (DLS) analysis showed comparable distributions of cumulant hydrodynamic radius (**Fig. 4f**), polydispersity index (**Fig. 4g**), and scattering intensity (**Fig. 4h**). Across all six attributes, no significant difference was detected between parental and refined sequences (two-sided Wilcoxon signed-rank test, all *p* > 0.05).

These results demonstrate that SynCodonLM-guided refinement systematically improves CHO titer, with improvement magnitude tracking the model’s likelihood signal, while preserving product identity (including native PTMs) and biophysical quality. More broadly, they show that clinically optimized therapeutic coding sequences remain evolvable despite extensive prior development, supporting iterative refinement as a practical strategy for engineering coding DNA independently of protein sequence.

### Mechanistic insight into differential expression

To probe the mechanistic basis of the observed titer improvements, we asked (i) whether SynCodonLM-guided refinement systematically shifted established codon-usage descriptors in directions heuristically associated with higher translational efficiency, and (ii) whether the SynCodonLM Evo score predicted per-construct titer improvements more effectively than classical single-metric heuristics.

We first evaluated the direction and significance of change for a broad panel of >23 established codon-usage descriptors (full list and references in Methods)^33^ computed on parental and refined sequence pairs. For each metric, the paired difference (Δ = refined − parental) was tested with a Wilcoxon signed-rank test, and each metric was annotated with a biologically expected direction for improved translational efficiency drawn from prior literature. Of the metrics with a defined directional expectation, the overwhelming majority shifted in the favorable direction (**Fig. 5a**), including CAI^16^, RSCU^34^, ENc^35–37^, GC content^38^, CpG and UpA O/E enrichment^39,40^, CBI^41^, and FOP^42^ (all *p* < 0.001); no directional descriptor shifted significantly in the opposing direction. Despite optimizing purely for codon-language-model likelihood, with no explicit metric-based objective, SynCodonLM refinement recovered the directional expectations of canonical codon-optimization heuristics across all descriptors at once.

**Fig. 5:**
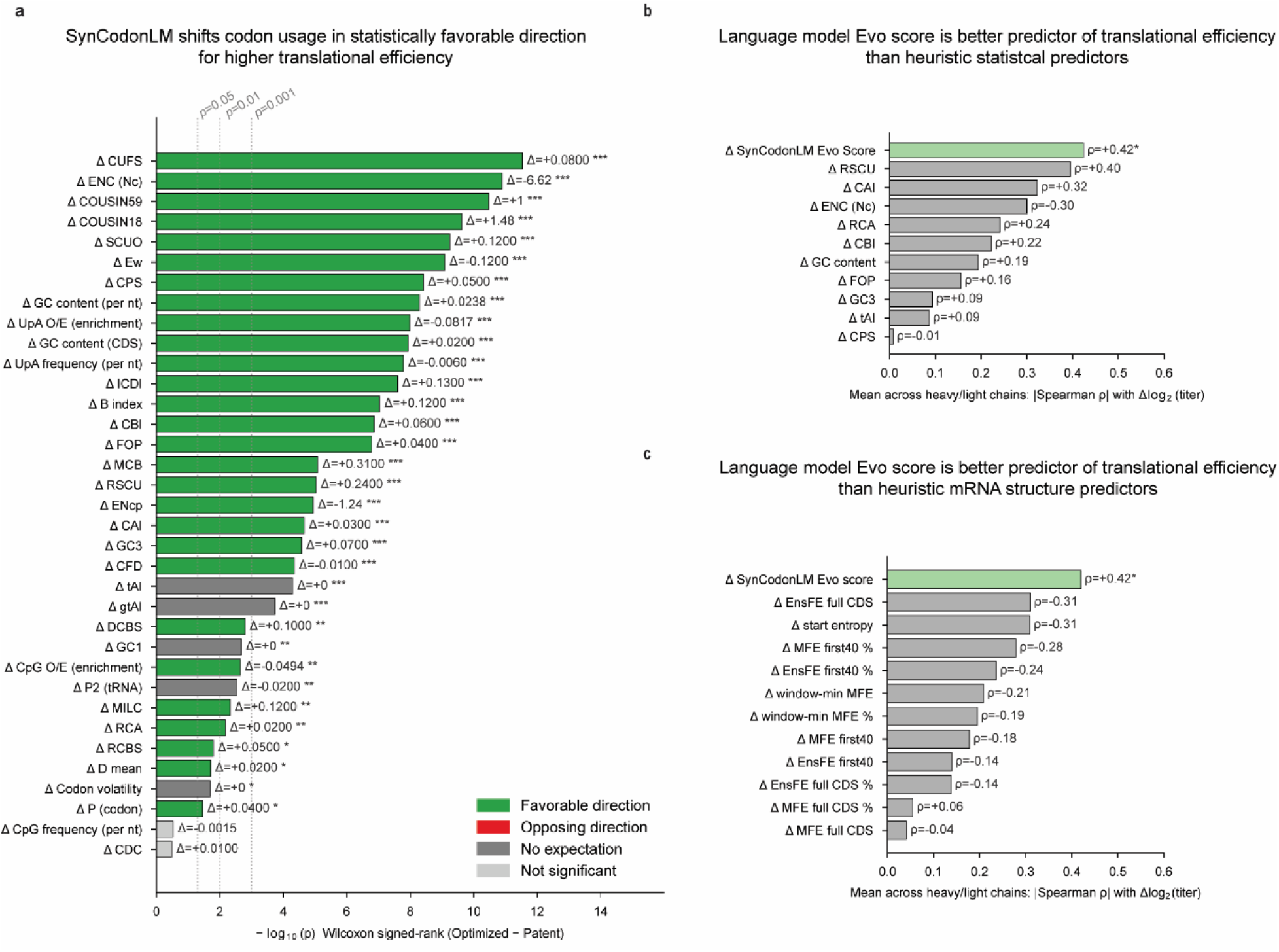
Codon language model likelihood outperforms classical codon-usage and mRNA-structure heuristics. (a) Paired Wilcoxon signed-rank comparison of established codon-usage descriptors on parental versus SynCodonLM-refined sequences (n = 23). The x-axis reports −log₁₀(*p*); dashed guides mark *p* = 0.05, 0.01, and 0.001. Bars are annotated with the paired difference (refined − parental) and colored by the expected direction for improved translational efficiency: green, favorable shift; red, opposing shift; dark gray, no defined expectation; light gray, not significant. \**p* < 0.05, \*\**p* < 0.01, \*\*\**p* < 0.001. (b) Mean absolute Spearman’s rank correlation coefficient (|ρ|, averaged across heavy and light chains) between per-descriptor change and Δlog₂(titer). Signed ρ values are annotated on each bar. The SynCodonLM Evo score (pastel green) is the strongest single predictor (ρ = +0.42, *p* < 0.05), exceeding all classical codon-usage heuristics (gray). (c) Analogous analysis using ViennaRNA-derived mRNA-structure descriptors. SynCodonLM Evo score again ranks first.

We next asked whether these classical descriptors were quantitatively predictive of the observed titer improvements. For each construct, we computed the mean absolute Spearman’s rank correlation across heavy and light chains between the per-metric change and the observed Δlog₂(titer) (**Fig. 5b).** The change in SynCodonLM Evo score was the strongest single predictor (ρ = +0.42, *p* < 0.05), outperforming every classical codon-usage descriptor evaluated (|ρ| ≤ 0.40; **Fig. 5b**). Notably, several widely used metrics (CAI, tAI^43^, GC3, FOP) showed weak correlations with translational efficiency, underscoring that individual heuristic descriptors capture only limited information about expression outcomes, even when they respond systematically in aggregate. A graph including all the statistical descriptors and their correlation with Δlog₂(titer) is available in **Supplementary Figure 6**. A nonparametric bootstrap (5,000 iterations; **Supplementary Fig. 7**) yielded broad confidence intervals for all descriptors, with substantial overlap among metrics, although the SynCodonLM Evo score retained the highest point estimate.

Because expression can also be shaped by mRNA secondary-structure features around the ribosome-loading region, a signal established by Kudla et al. in *E. coli*^44^, we performed a similar analysis using a panel of ViennaRNA-derived^45^ mRNA-structure descriptors (**Fig. 5c**; full descriptor definitions in Methods). The SynCodonLM Evo score again emerged as the strongest predictor of Δlog₂(titer) (ρ = +0.42, *p* < 0.05). Among the mRNA-structure descriptors, ensemble free-energy and sequence-entropy metrics showed the strongest associations with expression (|ρ| up to 0.31), whereas the canonical 5′ folding-window descriptors like that of Kudla et al., exhibited weaker correlations (|ρ| ≤ 0.28). Correlation strengths varied depending on whether the signal peptide was included in the folding calculation (**Supplementary Fig. 8**).

We further asked whether SynCodonLM’s suggested substitutions were concentrated at specific positions along the CDS, for example enriched near the 5′ ribosome-loading region where structure-mediated effects on translation initiation and early elongation are strongest^46,47^. We observed a clear positional bias, with edits enriched toward the 5′ end of the coding sequence and declining toward the 3′ end (**Supplementary Fig. 9**). To test the classical hypothesis that synonymous codon choice is coordinated with local protein structure^5,48^, we then examined editing rates relative to secondary-structure elements while controlling for codon degeneracy throughout each sequence. Codons at the start (N-cap) of β-sheet strands were significantly less likely to be edited than expected, consistent with the idea that codon usage at some structural boundaries is under selective constraint (**Supplementary Fig. 10**).

These results point to a non-obvious conclusion: therapeutic coding sequences retain accessible optimization potential beyond what can be captured by any single classical codon-usage or mRNA-structure heuristic. Although our iterative refinement strategy systematically moves sequences in the favorable direction of nearly all canonical codon-usage descriptors, no single descriptor predicts per-construct titer gains, whereas codon-language-model likelihood does. This suggests that residual coding-sequence optimization potential is distributed across many context-dependent features that classical heuristics capture only in isolation.

## Discussion

We reframed codon optimization as an iterative, model-guided directed-evolution process operating strictly within synonymous codon space. Applied to the patented coding DNA of 23 clinically deployed antibody-based therapeutics, sequences already refined through years of industrial development, SynCodonLM improved CHO titer in 17 of 23 cases (∼1.34-fold change on average) with no significant degradation of monomer content, colloidal behavior or post-translational modification. The central result is that clinically validated coding sequences remain evolvable in synonymous codon space.

This parallels modern protein engineering, where protein language models improve even highly mature antibodies through small numbers of localized edits^23^; our results extend that logic to the synonymous layer of the genetic code. Practically, this reframes codon-optimized constructs not as finished artifacts locked in at DNA synthesis, but as starting points that remain open to further model-guided refinement, where small synonymous edits accumulate into meaningful expression gains without altering the encoded protein.

The mechanistic analysis in **Fig. 5** clarifies why language-model-guided refinement outperforms static, single-metric heuristics. In aggregate, SynCodonLM shifted nearly every classical codon-usage descriptor (CAI, RSCU, CBI, FOP, ENc, GC content, and CpG and UpA enrichment) toward values associated with efficient translation, despite being trained only on a synonym-constrained masked-language-modeling objective, with no explicit knowledge of any of these metrics. This convergence indicates that codon-language-model likelihood implicitly integrates the same statistical regularities that decades of classical codon-optimization work have each captured in isolation. Consistent with this, the SynCodonLM Evo score showed the strongest per-construct association with relative expression (ρ = 0.42, *p* < 0.05), with the classical and mRNA-structure descriptors tested here reaching comparable but somewhat weaker associations (|ρ| ≤ 0.40). Together, these observations suggest that the advantage of a codon language model is compositional: rather than relying on any single heuristic, it captures a distributed, context-dependent combination of features that each classical descriptor represents only partially.

That codon-level likelihood improves expression is consistent with our finding that it also encodes genuine evolutionary signal. On an Influenza A H1N1 benchmark spanning nucleoprotein, hemagglutinin, and neuraminidase, SynCodonLM Evo scores increased significantly with time (r = 0.65, 0.79, 0.41; all *p* < 1e-3), whereas ESM-2^49^ scores on matched translated sequences showed the opposite trend. This grounds the refinement results: the same likelihood that tracks real viral evolution is what drives the observed titer gains.

Several limitations warrant acknowledgment. First, our data derive from small-scale transient CHO cultures; although transient titers have been shown in previous studies to preserve rank-order in stable clones^50^, translation to stable-pool or clonal production settings remains to be tested. Second, refinement was capped at 10% of codon positions per construct, leaving open whether deeper traversal or non-greedy acquisition strategies would yield further gains or introduce product-quality trade-offs. Third, our analytical package (intact LC-MS, SEC, DLS) detected no significant shifts in molecular weight, glycoform envelope, monomer content, aggregation, or colloidal behavior, but did not include manufacturing-depth characterization (released N-glycan mapping, icIEF/CEX charge-variant analysis, or site-specific peptide mapping). Fourth, the mechanistic analysis in **Fig. 5** is correlative and panel-level; it does not isolate which mechanistic axis is causal in any given molecule. Fifth, the viral evolution benchmark is subject to known geographic and temporal sampling biases and should be interpreted as evidence that codon-level likelihood tracks evolutionary progression, not as a claim that codon-level fitness causally shapes viral evolution.

Looking forward, the framework introduced here suggests several natural extensions. Iterative synonymous refinement is model-agnostic at the framework level: as codon language models continue to improve, through larger context windows, multi-species conditioning, or objectives explicitly coupled to downstream fitness signals, the same directed-evolution loop can be plugged in without changing its structure. Coupling refinement to closed-loop experimental feedback, in which measured titers or product-quality attributes inform the next round of edits, is a particularly natural next step and would convert the framework from a single-shot in silico refinement into a genuine active-learning cycle in coding DNA space. Beyond therapeutic antibodies, the same approach should generalize to any recombinantly expressed protein where the coding sequence is a controllable design axis, including vaccine antigens, gene-therapy payloads, and other biologics classes where synonymous choice has been shown to matter but has historically been treated as static.

More broadly, our results argue that the boundary between protein-level and codon-level engineering deserves to be revisited. Protein language models have unlocked iterative, evolutionarily plausible improvement at the amino-acid level^23^; codon language models now appear to offer an analogous capability at the synonymous layer, without any of the regulatory or immunogenicity considerations that accompany changes to the protein sequence itself. Treating already-optimized coding sequences as high-fitness starting points for further iterative refinement, rather than as design endpoints, offers a simple, low-risk, and experimentally validated route to improving clinical biologics, and reframes the coding layer of the genetic code as an actively engineerable, not merely selectable, design space.

## Methods

### High-fitness coding sequence baseline (clinical-stage reference set)

A high-confidence benchmark of pre-optimized antibody coding DNA sequences was curated by starting from Thera-SAbDab^51^, which tracks WHO-recognized antibody- and nanobody-derived therapeutics and provides associated sequence and metadata. As a first pass, each therapeutic’s variable (V) region was used to query the patent nucleotide collection with NCBI BLAST^52^ (blastn against nt), retaining only patent-disclosed coding DNA sequences that returned 100% identity across the V region and therefore encoded the same V region as the clinical molecule. For each therapeutic entry, primary patent documents and sequence listings were mined using automated document-review workflows to determine whether full-length coding DNA sequences were disclosed for each expressed chain of the final drug molecule and whether those sequences were described in a CHO expression context. Candidate chains were then cross-mapped to publicly available substance registration resources in the Global Substance Registration System (GSRS)^53^ to corroborate identity and chain assignment through sequence-based checks aligned with regulated substance definitions and sequence search capabilities. The resulting collection served as a high-fitness baseline of coding sequences that were plausibly already optimized in an industrial therapeutic development context and therefore provided an empirically grounded starting point for in silico refinement.

In total, 23 clinical antibody-based therapeutics were gathered (comprising 22 distinct therapeutics corresponding to 23 expressed constructs, as the amivantamab bispecific was produced and analyzed as its two half-antibodies), including: monoclonal antibodies spanning IgG1, IgG2 and IgG4 heavy chain isotypes as well as both kappa and lambda light chain isotypes. The heavy-chain isotype distribution was dominated by IgG1 (n = 16) and IgG4 (n = 6), with a single IgG2 representative (fremanezumab), while light-chain use was almost exclusively kappa (n = 22) with lambda represented by a single chain (belimumab); isotype pairings therefore break down as IgG1κ (n = 15), IgG1λ (n = 1), IgG2κ (n = 1), and IgG4κ (n = 6). Nineteen chains were derived from approved medicines, two from Phase II candidates, and two from a Phase I/ candidate (**Supplementary Table 1**). Beyond conventional monospecific IgG monoclonal antibodies (n = 18), the reference set was deliberately enriched for architecturally complex, industrially relevant formats whose coding sequences must nonetheless be engineered for high-fitness expression in CHO: one antibody–drug conjugate (brentuximab vedotin, a chimeric IgG1κ site-conjugated on reduced interchain cysteines via a cathepsin-cleavable valine-citrulline linker to monomethyl auristatin E at a drug-to-antibody ratio of ≈ 4)^54^; two bispecific IgG molecules: amivantamab (a Genmab DuoBody IgG1κ target-bispecific against EGFR × MET, assembled from two half-antibodies bearing complementary F405L/K409R Fc mutations that undergo controlled Fab-arm exchange^55^ and produced in low-fucose CHO to enhance ADCC), marlotamig (a Regeneron Veloci-Bi IgG4 with a common kappa light chain and a "chimeric star-Fc" heavy-chain mutation on one arm that ablates Protein A binding to enforce heterodimer-selective purification^56^); and three antibody fusion proteins: melredableukin alfa (a non-targeting IgG1κ scaffold C-terminally fused to an engineered CD25-silent IL-2 variant, presented monovalently via knob-into-hole Fc pairing^57^), AGT-182 (a HIRMab-IDS fusion: an anti-human-insulin-receptor IgG1κ blood–brain-barrier shuttle with iduronate-2-sulfatase fused to the heavy-chain C-terminus for CNS enzyme delivery in MPS II), and vislarafusp alfa (an anti-HER2 IgG1κ with the SIRPα D1 domain appended to the light-chain N-terminus via a GS linker, acting as a locally-delivered CD47 decoy). Several molecules additionally carry defined Fc engineering, including the YTE (M252Y/S254T/T256E) FcRn-affinity mutation that extends serum half-life in nirsevimab^58^ and the IgG4 hinge-stabilizing S228P substitution that prevents natural IgG4 Fab-arm exchange present across the IgG4 mAbs^59^. Full per-molecule annotation, format assignments, targets, and the primary patent identifier used to source each coding sequence are provided in **Supplementary Table 1**.

For 5 of the 23 constructs, the coding sequence was obtained from a patent sequence listing assigned to an entity other than the originator of the marketed therapeutic; in each of these cases the disclosed sequence nonetheless encodes the corresponding therapeutic protein, consistent with the sequence-based identity checks described above.

In a small number of constructs, the patent-disclosed coding DNA did not translate to the exact clinical protein sequence at every position. In some of these cases the patent text identified the disclosed sequence as encoding the final therapeutic protein, while the deposited coding DNA translated to a small difference from that protein, consistent with isolated discrepancies within the sequence listings themselves. First, the belimumab and brentuximab coding sequences each carried a C-terminal lysine on the heavy chain that is absent from the final molecule listed in GSRS; C-terminal lysines are, however, routinely clipped from mammalian expression products and are therefore not expected to affect the expressed protein. The bimekizumab coding sequence contained a single-residue difference in the Fc region of its heavy chain at an atypical position. The amivantamab half-antibody 1 light-chain coding sequence was missing the N-terminal alanine. Lastly, the enlonstobart coding sequence encoded an IgG1 backbone rather than the IgG4 molecule listed in GSRS, although its V region and full light chain matched the final drug molecule. In each case the discrepancy was individually verified against the GSRS/INN reference and the coding sequence retained for downstream analysis.

### SynCodonLM inference details

SynCodonLM was loaded from the Version 2 model checkpoint on HuggingFace^24,60^ onto a Nvidia L40S GPU running CUDA 13.0 for all evaluations in this work. Environment versioning is available on the corresponding GitHub codebase released with this work.

### SynCodonLM-based evolutionary scoring and greedy synonymous directed evolution

Synonymous codon refinement was performed with SynCodonLM^24^, a codon language model trained under a biologically grounded constraint in which masked codons were predicted only from synonymous options consistent with the known amino-acid sequence, thereby disentangling codon-level patterns from protein-level semantics. Each coding sequence was represented as a sequence of codon tokens x = (*x*_1_,...., *x_L_*), where L denotes the number of codons. To quantify “evolutionary” compatibility of a complete sequence using a masked language model, a pseudo-log-likelihood (PLL) score was computed by iteratively masking one position at a time and accumulating the log probability assigned to the true token at that position given the remaining unmasked context. This single-mask PLL scoring paradigm is a standard approach for scoring sequences with masked language models in the absence of an explicit left-to-right factorization. Specifically, for position *i*, the model produced logits *z_i_*(ν) over vocabulary tokens ν ∈ V when *x_i_* was replaced by a mask token, and the per-position log probability of the true codon token was computed as

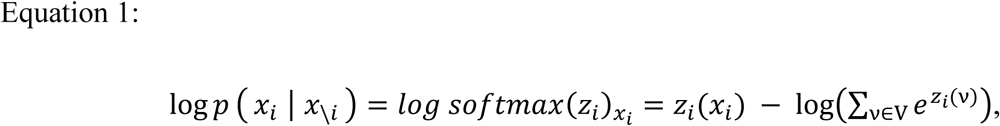

where *x*_\*i*_ denotes the masked sequence with all positions except *i* left intact. The sequence-level evolutionary score was reported as the mean PLL across codon positions (in nats),

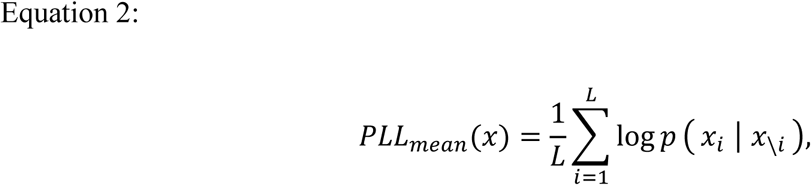

with higher values indicating greater model compatibility.

To propose synonymous edits, scoring at each position was restricted to the set of synonymous codons for the amino acid *a_i_* encoded at position *i*. Let S(*a_i_*) denote the synonymous codon set at that position. Scores were normalized over the synonym set through a synonym-restricted softmax, rather than the full-vocabulary softmax, such that

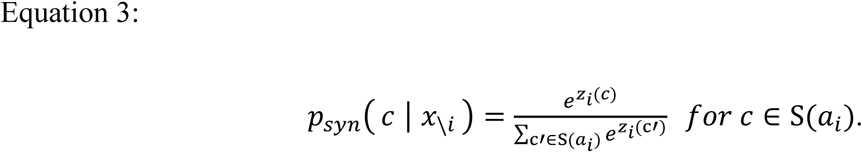

This ensured that positions with different synonym degeneracies were compared within their appropriate synonymous choice sets, mirroring the normalization implemented by restricting logits to the synonymous token IDs at each step. Greedy optimization proceeded by iteratively selecting a single synonymous substitution that maximized a head-to-head Δ-margin at one position. For a candidate position *i*, with current codon *c_curr_* = *x_i_* and best alternative synonym *c_best_* ∈ S(*a_i_*) \ (*c_curr_*), the step score was defined on the logit scale as

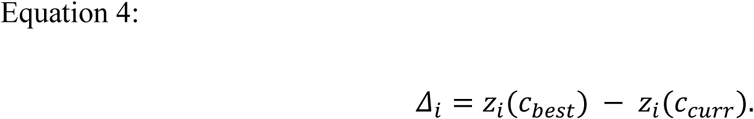

Because both terms share the same synonym-restricted normalization constant, *Δ_i_* was equivalently the difference in synonym-normalized log probabilities,

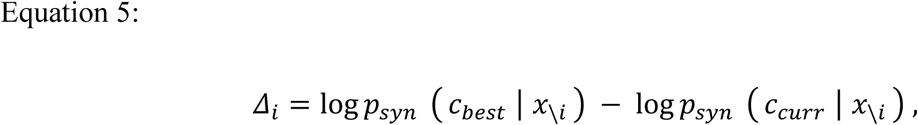

and the corresponding odds ratio reported during optimization was

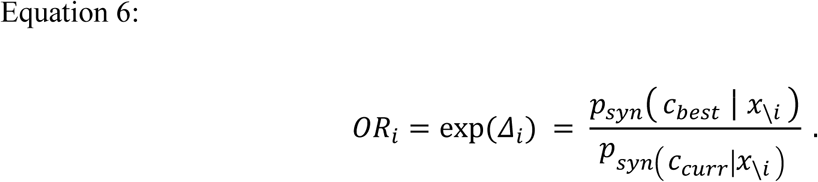

At each iteration, the single position and substitution with the largest admissible *Δ_i_* were applied, the sequence context was updated, and the process was repeated until no admissible Δ remained above threshold.

To constrain divergence from clinically validated parental sequences, the number of codon positions altered relative to the original sequence was capped at ⌊0.10L⌋ (10% of codon positions), with optional tightening of Δ thresholds once the cap was reached to avoid negligible micro-steps. Start and stop boundaries were preserved when configured (retaining an initiating ATG and terminal stop codon), and short-cycle oscillations were mitigated by immediate reversal checks, state-hash cycle detection, and a small tabu memory over recently edited positions. Species conditioning was applied via a species token type, enabling host-contextualized scoring under the model’s token-type mechanism when supported.

### Cloning, gene synthesis and construct assembly

Parental and SynCodonLM-refined coding sequences were synthesized as single linear DNA fragments (Twist Bioscience) with flanking BsaI recognition sites compatible with Type IIS Golden Gate assembly. Each construct was assembled into a pTT5-based mammalian expression vector^31^ containing a fixed signal peptide and stop codon using a BsaI-based Golden Gate cloning kit (New England Biolabs) in a single one-pot reaction per construct, following the manufacturer’s recommended thermocycling protocol. Assembly reactions were transformed into chemically competent E. coli, plated on selective medium, and individual colonies were picked for downstream screening.

Colonies were screened by colony PCR using primers flanking the insertion site, and PCR amplicons spanning the full open reading frame (ORF) were verified by Illumina MiSeq short-read sequencing. Each construct was considered sequence-confirmed only upon full-ORF concordance with the intended parental or refined coding sequence, including all synonymous positions edited by SynCodonLM. Sequencing was carried out iteratively until a colony passing full-ORF verification was recovered for every construct.

Sequence-confirmed colonies were then scaled up in selective liquid culture, and endotoxin-reduced plasmid DNA was prepared using the QIAGEN Plasmid Plus midiprep kit (QIAGEN) according to the manufacturer’s protocol. Purified plasmid preparations were quantified by UV absorbance (A₂₆₀), assessed for purity by A₂₆₀/A₂₈₀ ratio, and used directly for transient CHO transfections.

### CHO expression and antibody purification

Transient expression was performed in CHO-3E7 cells^32^, a platform described to enhance recombinant protein production when paired with oriP-bearing pTT vectors and optimized CMV-based expression cassettes.

Cells were maintained in Irvine Scientific BalanCD CHO medium and seeded at a density of 4 × 10^6^ cells/mL with viability >99% at the time of transfection. Transfections were carried out using TransIT-PRO (Mirus Bio) in OptiPRO SFM (Thermo Fisher Scientific) according to the manufacturer’s recommendations. Heavy- and light-chain expression plasmids were co-transfected at a 1:2 mass ratio (heavy:light), a commonly used configuration intended to promote efficient antibody assembly and secretion^61^. Approximately 16–24 hours post-transfection, cultures were supplemented with CHO Feed B (Irvine Scientific) and Anti-Clumping Agent (Thermo Fisher Scientific) to support cell health and productivity. Temperature was subsequently shifted from 37°C to 30 °C to enhance protein expression and stability during the production phase. Cultures grew for a total of 6 days in a culture volume of 4mL within 24-well plates.

Antibodies were purified from culture supernatants using an AssayMap Bravo platform (Agilent Technologies) equipped with Protein A affinity tips (Agilent Technologies). Purified antibodies were eluted and formulated into 75 mM citrate, 250 mM sodium phosphate, pH 5.4.

### Antibody titer quantification using Protein A affinity chromatograph on supernatant

The titer determination experiments were performed using an Agilent Bio-Monolith rProtein A column (4.95 × 5.2 mm; Agilent Technologies) on an Agilent 1290 Infinity II Bio LC system. The column temperature was maintained at 25°C. Detection was carried out using a fluorescence detector set at an excitation wavelength of 280 nm and an emission wavelength of 350 nm. Prior to analysis, cell culture supernatants were subjected to centrifugation at 3200 × g for 20 min, followed by filtration through a 0.2 µm filter plate. Samples were subsequently stored at 4 °C in the autosampler, and an injection volume of 20 µL was used for each run.

Mobile phase A comprised 50 mM sodium phosphate at pH 7.4, and mobile phase B was 100 mM citric acid at pH 2.6, both prepared in Milli-Q water using 1M stock solutions. Chromatographic separation was performed at a flow rate of 1 mL/min using the following gradient: initial equilibration with 100% mobile phase A for 0.8 min, followed by a linear gradient from 0% to 100% mobile phase B from 0.8 to 2.2 min to elute bound antibodies. The column was then re-equilibrated with 100% mobile phase A from 2.2 until the end of a 4.0-min run time. Data acquisition and analysis were performed using Agilent OpenLab software. Antibody titers were quantified by peak area integration against a Rituximab standard curve ranging from 10 to 1000 μg/ml.

### Mass spectrometry analyses

Detailed LC-MS parameters are provided in **Supplementary Table 2**. Briefly, intact and subunit level analyses were performed using a 6545XT Q-ToF interfaced with a 1290 Infinity II UHPLC (Agilent Technologies, Santa Clara, CA). Ballistic gradient conditions were used to desalt samples over a reversed-phase column (Poroshell SB 300 C3, 1.0 x 30 mm, 5 µm, 300Å, Agilent Technologies, Santa Clara, CA) prior to introduction into the mass spectrometer. Raw spectra were extracted and deconvoluted using MassHunter v11 software to enable (1) verification of predicted molecular weight values for each molecule and (2) pairwise comparison of the post-translational modifications within each molecule pair (parental and refined) at both the complexed (intact) and reduced subunit levels.

### Codon-usage descriptor calculations

Codon usage descriptors were calculated using the GenScript Rare Codon Analysis Tool^33^, utilizing the NCBI Genome Resource^62^ reference source, for *Cricetulus griseus* (CHO). Descriptors were computed on the mature coding sequence only; the fixed signal peptide and stop codon were excluded from all calculations, as these positions were held constant across parental and refined pairs and were not subject to synonymous refinement. Metrics computed included CAI^16^, tAI^43^, gtAI^63^, RSCU^34^, RCA^64^, CBI^41^, FOP^42^, CFD^65^, ENc/ENcp^35–37^, GC/GC1/GC3 content^38^, CpG and UpA dinucleotide statistics (frequency and observed/expected enrichment)^39,40^, CPS/CUFS^66–68^, COUSIN18/COUSIN59^69^, ICDI^70^, MCB^71^, MILC^72^, DCBS^73^, RCBS^74^, SCUO^75,76^, B index^77^, D mean^78^, Ew^79^, codon volatility^80^, P2(tRNA)^81^, P(codon)^82^, and CDC^83^. Biologically expected directionality for improved translational efficiency (used for color coding in **Fig. 5a**) was assigned from prior literature, metrics without an established directional expectation were annotated accordingly.

### mRNA secondary-structure analysis

mRNA secondary structure and associated thermodynamic descriptors were computed with the ViennaRNA package (RNAfold, v2.7.0)^45^ via its Python bindings, at 37 °C under default Turner 2004 thermodynamic parameters. DNA sequences were transcribed to RNA prior to folding, and all sequences were analyzed in both "with signal peptide" (wSP) and "without signal peptide" (noSP) configurations; the signal peptide and stop codon were held constant across all parental and refined pairs. For each sequence, minimum free energy (MFE) and partition-function ensemble free energy (EnsFE) were computed on the full coding sequence, and a sliding-window minimum MFE was calculated by scanning a 60-nt window in single-nucleotide steps along the full CDS and retaining the lowest window MFE together with its position; the 60-nt window size was chosen to capture typical mRNA hairpin/stem-loop length scales while preserving positional resolution across the transcript. The same MFE and EnsFE calculations were also performed on the 5′ prefix corresponding to the first 40 codons (120 nt) to characterize structure in the ribosome-loading region. To summarize ensemble-averaged structural content near the start codon, positional pairing probabilities were extracted from the partition-function base-pair probability matrix, converted to a binary paired/unpaired positional entropy, and averaged across the first 30 nucleotides to yield a scalar "start entropy" descriptor. Descriptors were computed for every expressed chain and averaged to yield per-molecule values, and paired differences (Δ = refined − parental) were used for all downstream statistical comparisons. Values reported in **Fig. 5c** correspond to the mature coding sequence without the signal peptide; the analogous wSP analysis is provided in **Supplementary Fig. 8** and yielded qualitatively concordant conclusions.

### Protein structure analysis

Three-dimensional structures were predicted for each expressed antibody chain with ESMFold^49^ (ESMFold2-Fast) on Tamarind Bio using default settings, folding each heavy and light chain independently as a monomer from its full-length sequence. Per-residue secondary structure and solvent accessibility were assigned with DSSP via Biopython^84^. DSSP’s eight states were collapsed to three (helix, strand, loop), relative solvent accessibility (rel_asa) was computed by normalizing each residue’s accessible surface area to its maximum reference value^85^, and residues below 0.20 were classified as buried. Backbone φ/ψ angles were used to assign α-helical and β-sheet Ramachandran basins, and per-residue pLDDT was retained as a confidence measure. For each contiguous helix and strand, we defined the first two residues as the N-cap (start), the last two as the C-cap (end), and the one to two preceding coil residues as the pre-element flank. Predicted structures were aligned to the coding sequence by translating the parental CDS and matching it to the modeled sequence, so each codon’s edit status could be mapped to its residue’s features; only chains with at least 20 aligned codons were retained.

To test whether editing was associated with any structural context independently of codon redundancy, each feature was fit in its own logistic regression predicting per-codon edit status, controlling for codon degeneracy (edited ∼ feature + degeneracy). Binary features were modeled as context versus all other residues, and continuous features (rel_asa, pLDDT, normalized 5′→3′ position) were standardized and reported per standard deviation. Standard errors were clustered by chain, effects are reported as odds ratios with 95% confidence intervals, and p-values were corrected across the panel using the Benjamini-Hochberg false discovery rate^86^ (*q* < 0.05 significant; **Supplementary Fig. 10**). Analyses used Biopython, pandas, NumPy, SciPy, and statsmodels in the same versions listed above.

### Statistical analysis

All statistical analyses were performed in Python 3.12. Analyses supporting **Figs. 4** and **5** were run under Python 3.12.0 using SciPy v1.13.0, NumPy v1.26.4, pandas v2.2.3, and statsmodels v0.14.2, while the viral evolution analyses supporting **Fig. 2** were run under Python 3.12.2 in a separate high-performance computing environment using SciPy v1.15.1, NumPy v2.0.1, pandas v2.2.3, statsmodels v0.14.5, PyTorch v2.7.1 (CUDA 12.6), Transformers v4.48.3, and scikit-learn v1.6.1. Random seeds were fixed for all resampling procedures to ensure reproducibility. For the titer analysis in **Fig. 4**, per-construct log₂-fold change was computed as log₂(mean refined titer / mean parental titer) across biological triplicates, and panel-wide comparison of refined versus parental log₂-titer was assessed using both a two-sided paired t-test and a two-sided Wilcoxon signed-rank test. The relationship between the change in mean chain-level SynCodonLM Evo score (ΔEvo = refined − parental, averaged across all expressed chains of a molecule) and the observed Δlog₂(titer) was quantified by Spearman’s rank correlation across n = 23 molecules. For the mechanistic analyses in **Fig. 5**, paired differences (Δ = refined − parental) for each codon-usage descriptor and each ViennaRNA-derived mRNA-structure descriptor were tested using a two-sided Wilcoxon signed-rank test, and per-descriptor correlations with Δlog₂(titer) were computed as the mean absolute Spearman’s ρ across heavy- and light-chain-level computations, with signed ρ values reported alongside. To assess robustness of the per-descriptor rankings to individual influential molecules, a nonparametric case-resampled bootstrap was performed with 5,000 iterations using NumPy’s PCG64 generator (seed = 9); on each iteration, molecules were sampled with replacement (n = 23) and Spearman’s correlations between each descriptor and Δlog₂(titer) were recomputed, with iterations discarded when either variable lacked variance in the resample. Two-sided 95% confidence intervals were derived using the percentile method (2.5th and 97.5th percentiles across valid resamples), and a minimum of 100 valid resamples was required for a CI to be reported. For the viral evolution analyses in **Fig. 2**, Spearman’s and Pearson correlations between per-sequence Evo score and sampling year were computed independently for each Influenza A H1N1 protein (nucleoprotein, hemagglutinin, neuraminidase), with R² reported as the square of the Pearson r. Because sample sizes were large (N = 855–6,710), p-values below the numerical precision floor of the underlying SciPy implementation are reported as *p* < 1 × 10^−3^.

### Analytical-SEC to measure aggregation and protein purity

Antibodies were analyzed by analytical size exclusion chromatography (aSEC) to assess monomer purity and aggregation. Analyses were performed using an Agilent 1290 Infinity II system equipped with an ACQUITY UPLC Protein BEH SEC column (200 Å, 1.7 µm, 4.6 × 150 mm; Waters Corp., Milford, MA).

The separation was conducted at a flow rate of 0.5 mL/min over a 5 min runtime at 37 °C. Samples were eluted using a mobile phase consisting of 50 mM sodium phosphate, 200 mM L-arginine hydrochloride, and 0.05% (w/v) sodium azide at pH 6.8. The mobile phase was prepared fresh, filtered, and degassed prior to use.

Elution was monitored by UV absorbance at 280 nm. Monomer and aggregate species, including low molecular weight (LMW) and high molecular weight (HMW) components, were quantified based on peak area integration via Agilent OpenLab software.

### Dynamic light scattering to provide further measurements on aggregation and protein purity

Dynamic light scattering (DLS) was performed to further evaluate protein aggregation and sample homogeneity. Measurements were conducted using a Prometheus Panta instrument (NanoTemper Technologies, Munich, Germany). Samples were analyzed under native conditions without dilution unless otherwise specified.

Measurements were carried out at 25 °C with automatic laser intensity optimization. The cumulant radius, polydispersity index, and scattering intensity were determined from the scattered light intensity using instrument-provided analysis software.

Size distributions were calculated based on intensity-weighted analysis to detect the presence of monomeric species and higher-order aggregates. Data were used to assess sample purity, aggregation propensity, and colloidal stability, complementing chromatographic analysis obtained by aSEC.

### Viral evolution dataset curation

We curated a viral evolution benchmark from the NCBI Virus database ^87^ by querying for Alphainfluenzavirus influenzae sequences isolated from human hosts and corresponding to genome segment 5, which encodes nucleoprotein. Coding sequences and associated metadata were downloaded and restricted to H1N1 isolates. Sequences were then filtered to retain only complete, in-frame coding DNA sequences: records containing non-ATGC characters, internal stop codons, lengths not divisible by three, or missing terminal stop codons were excluded.

To enable a controlled comparison between codon- and protein-level language models, coding sequences were collapsed by translated amino-acid sequence. Specifically, when multiple synonymous coding sequences encoded the same nucleoprotein sequence, only the earliest available coding-sequence representative for that protein sequence was retained. This deduplication strategy preserved a one-to-one mapping between each protein sequence and a single representative coding sequence, ensuring that SynCodonLM and ESM-2 were evaluated on matched evolutionary sequence instances rather than on unequal codon-level redundancy.

### Viral Evolution Evo Score Calculations

Evolutionary compatibility scores were computed separately for matched coding and protein sequences using SynCodonLM and ESM-2, respectively. SynCodonLM was applied to the full viral coding sequence, including the initiating ATG and terminal stop codon. ESM-2 was applied to the corresponding translated protein sequence from the initiating methionine through the final amino acid, excluding the stop codon because stop symbols are not represented as standard amino-acid tokens in the protein model vocabulary. ESM-2 is a transformer-based protein language model distributed as part of the Evolutionary Scale Modeling family of pretrained protein models, and ESM-2-style implementations support masked-token prediction and per-token logits suitable for sequence scoring.

For each model, an Evo score was calculated using a single-position masked pseudo-log-likelihood framework. For a sequence x = (*x*_1_,...., *x_L_*), each position *i* was masked independently, and the model-assigned log probability of the observed token *x_i_*was computed conditional on the remaining unmasked sequence context *x*_\*i*_. The sequence-level Evo score was then defined as the mean pseudo-log-likelihood across all scored positions:

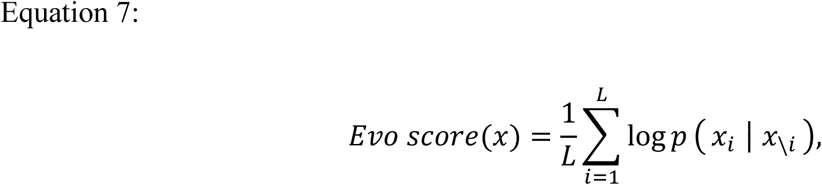

where *x_i_* denotes either a codon token for SynCodonLM or an amino-acid token for ESM-2. Higher Evo scores indicate greater compatibility between the observed sequence and the evolutionary/statistical constraints learned by the corresponding language model. Because SynCodonLM and ESM-2 operate over different token spaces and were trained under different modeling objectives, Evo scores were interpreted within each model rather than as directly interchangeable absolute quantities across models. Comparisons between codon- and protein-level scoring were therefore based on relative trends across matched viral sequence instances.

### AI Usage

Microsoft Copilot was used to assist with drafting portions of the manuscript and with code development. All content was reviewed, edited, and validated by the authors.

## Code Availability

A repository to reproduce the work in this article, as well as utilize the model for future purposes, is available at https://github.com/Boehringer-Ingelheim/Directed-Evolution-in-Codon-Space. The software is fully open source and built for ease of use and implementation.

## Data Availability

Data is available attached to this article in **Supplemental Data**.

## Acknowledgements

We thank Boehringer Ingelheim for providing the high-performance computing resources as well as wet lab, high throughput molecular biology facilities utilized in this work. J.H. gratefully acknowledges funding support from S.C. through his institute professor’s endowed chair, as well as additional support from Boehringer Ingelheim.

