## Supplementary Information for "Directed Evolution in Codon Space"

| INN | Target | Molecule Format | Patent Link |
|---|---|---|---|
| Amivantamab | EGFR × MET (c-Met) | Bispecific IgG1, κ/κ (DuoBody) | https://patents.google.com/patent/US9695242B2/en |
| Belimumab | BAFF (TNFSF13B) | IgG1, λ monoclonal antibody | https://patents.google.com/patent/EP3059319A1/en |
| Bimekizumab | IL-17A (and IL-17F) | IgG1, κ monoclonal antibody | https://patents.google.com/patent/US9034600B2/en |
| Brentuximab (vedotin) | CD30 (TNFRSF8) | IgG1, κ antibody–drug conjugate (ADC) | https://patents.google.com/patent/US10864277B2/en |
| Burosumab | FGF23 | IgG1, κ monoclonal antibody | https://patents.google.com/patent/US9290569B2/en |
| Canakinumab | IL-1β | IgG1, κ monoclonal antibody | https://patents.google.com/patent/WO2018229612A1/en |
| Concizumab | TFPI | IgG4, κ monoclonal antibody | https://patents.google.com/patent/US9795674B2/en |
| Daclizumab | CD25 (IL-2Rα) | IgG1, κ monoclonal antibody | https://patents.google.com/patent/US9815903B2/en |
| Donanemab | Aβ (pyroglutamate-modified amyloid plaques) | IgG1, κ monoclonal antibody | https://patents.google.com/patent/US8961972B2/en |
| Dostarlimab | PD-1 | IgG4, κ monoclonal antibody | https://patents.google.com/patent/US11155624B2/en |
| Enlonstobart (Isotype variant) | PD-1 | IgG1, κ monoclonal antibody (IgG4 in clinic) | https://patents.google.com/patent/US10774144B2/en |
| Fremanezumab | CGRP (α and β) | IgG2, κ monoclonal antibody | https://patents.google.com/patent/US9884907B2/en |

| | | | |
|---|---|---|---|
| Galcanezumab | CGRP (α and β) | IgG4, κ monoclonal antibody | https://patents.google.com/patent/US20230159628A1/en |
| Infliximab | TNF-α | IgG1, κ monoclonal antibody | https://patents.google.com/patent/US10066019B2/en |
| Marlotamig | EGFR × CD28 | Bispecific IgG4, common-κ light chain (Regeneron Veloci-Bi™) | https://patents.google.com/patent/US20200299388A1/en |
| Melredableukin (alfa) | Non-binding IgG1 (IL-2Rβγ agonist payload) | IgG1 Fc-fused IL-2 mutein (immunocytokine) | https://patents.google.com/patent/US11098099B2/en |
| Mirikizumab | IL-23 p19 | IgG4, κ monoclonal antibody | https://patents.google.com/patent/US9023358B2/en |
| Nirsevimab | RSV F protein (prefusion) | IgG1, κ monoclonal antibody (YTE half-life extended) | https://patents.google.com/patent/US10774133B2/en |
| HIRMab-IDS (AGT-182) (alfa) | INSR / HIR (BBB shuttle) + IDS enzyme | IgG1, κ antibody–enzyme fusion (BBB shuttle) | https://patents.google.com/patent/US8834874B2/en |
| Serplulimab | PD-1 | IgG4, κ monoclonal antibody | https://patents.google.com/patent/US11028173B2/en |
| Vedolizumab | α4β7 integrin | IgG1, κ monoclonal antibody | https://patents.google.com/patent/US20220267448A1/en |
| Vislarafusp (alfa) | HER2 × CD47 (SIRPα decoy) | IgG1, κ SIRPα-decoy fusion antibody | https://patents.google.com/patent/US11453724B2/en |

**Supplementary Table 1: Evaluation dataset properties**

| INN | Mass Spec Conditions |
|---|---|
| Amivantamab (Both Halves) | 1. Reduced<br>2. Reduced + Deglycosylation |
| Belimumab | 1. N/A<br>2. Reduced + Deglycosylation |

| | |
|---|---|
| Bimekizumab | 1. Reduced<br>2. Reduced + Deglycosylation |
| Brentuximab (vedotin) | 1. Reduced<br>2. Reduced + Deglycosylation |
| Burosumab | 1. Reduced<br>2. Reduced + Deglycosylation |
| Canakinumab | 1. Reduced<br>2. Reduced + Deglycosylation |
| Concizumab | 1. Reduced<br>2. Reduced + Deglycosylation |
| Daclizumab | 1. Reduced<br>2. Reduced + Deglycosylation |
| Donanemab | 1. Reduced<br>2. Reduced + Deglycosylation |
| Dostarlimab | 1. Reduced<br>2. Reduced + Deglycosylation |
| Enlonstobart | 1. Reduced<br>2. Reduced + Deglycosylation |
| Fremanezumab | 1. Reduced<br>2. Reduced + Deglycosylation |
| Galcanezumab | 1. Reduced<br>2. Reduced + Deglycosylation |
| Infliximab | 1. Reduced<br>2. Reduced + Deglycosylation |
| Marlotamig | 1. Reduced<br>2. Reduced + Deglycosylation |
| Melredableukin (alfa) | 1. Reduced<br>2. Reduced + Deglycosylation |
| Mirikizumab | 1. Reduced<br>2. Reduced + Deglycosylation |
| Nirsevimab | 1. Reduced<br>2. Reduced + Deglycosylation |
| HIRMab (AGT-182) (alfa) | 1. Reduced (could not analyze heavy chain due to 8 glycoforms on fusion protein) |

| | |
|---|---|
| Serplulimab | 1. Reduced<br>2. Reduced + Deglycosylation |
| Vedolizumab | 1. Reduced<br>2. Reduced + Deglycosylation |
| Vislarafusp (alfa) | 1. Reduced<br>2. Reduced + Deglycosylation |

**Supplementary Table 2: Mass spec conditions performed on evaluation dataset**

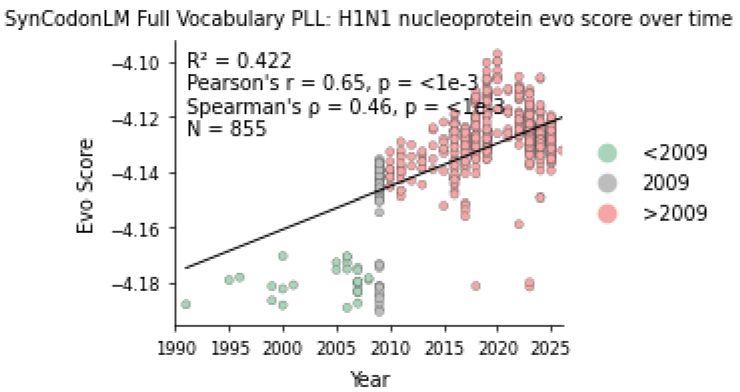


**Supplementary Figure 1: Full-vocabulary codon likelihood recovers the temporal signal for H1N1 nucleoprotein.** Non-synonym-constrained SynCodonLM pseudo-log-likelihood versus sampling year (Pearson r = 0.65, $p < 1 \times 10^{-3}$; Spearman ρ = 0.46, $p < 1 \times 10^{-3}$; N = 855), closely matching the synonym-constrained score in **Fig. 2a**.

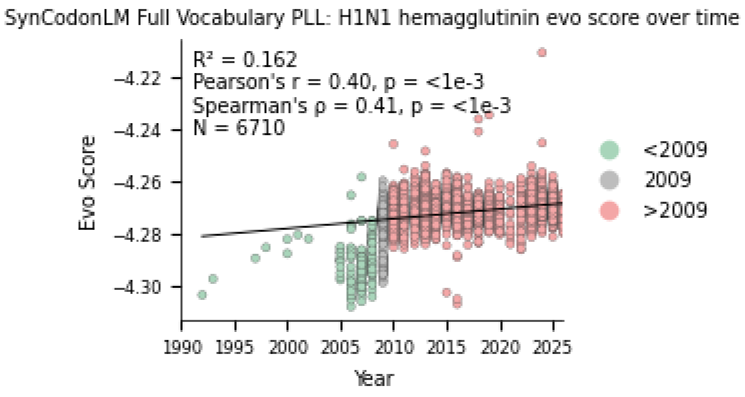


**Supplementary Figure 2: Full-vocabulary codon likelihood retains a positive temporal signal for H1N1 hemagglutinin.** (Pearson r = 0.40, $p < 1 \times 10^{-3}$; Spearman ρ = 0.41, $p < 1 \times 10^{-3}$; N = 6,710), weaker than the synonym-constrained score in **Fig. 2c** but consistent in direction.

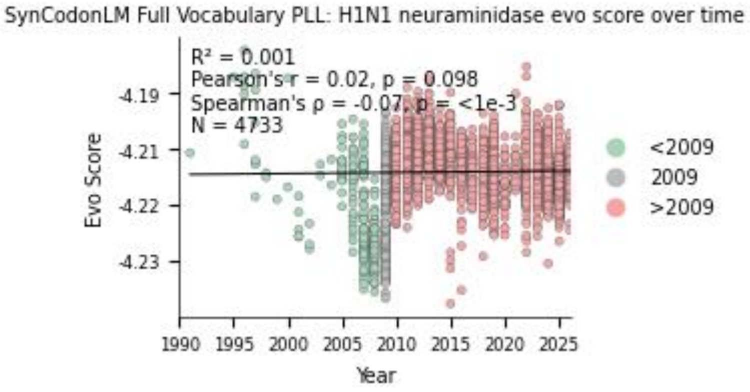


**Supplementary Figure 3: For H1N1 neuraminidase, the temporal signal is lost under full-vocabulary scoring** (Pearson r = 0.02, $p = 0.098$; Spearman ρ = −0.07; N = 4,733), whereas synonym-constrained scoring retains it (**Fig. 2e**, r = 0.41). This shows that synonym-constrained scoring, rather than full-vocabulary likelihood, is required to recover the neuraminidase signal, underscoring the value of the synonym constraint.

.

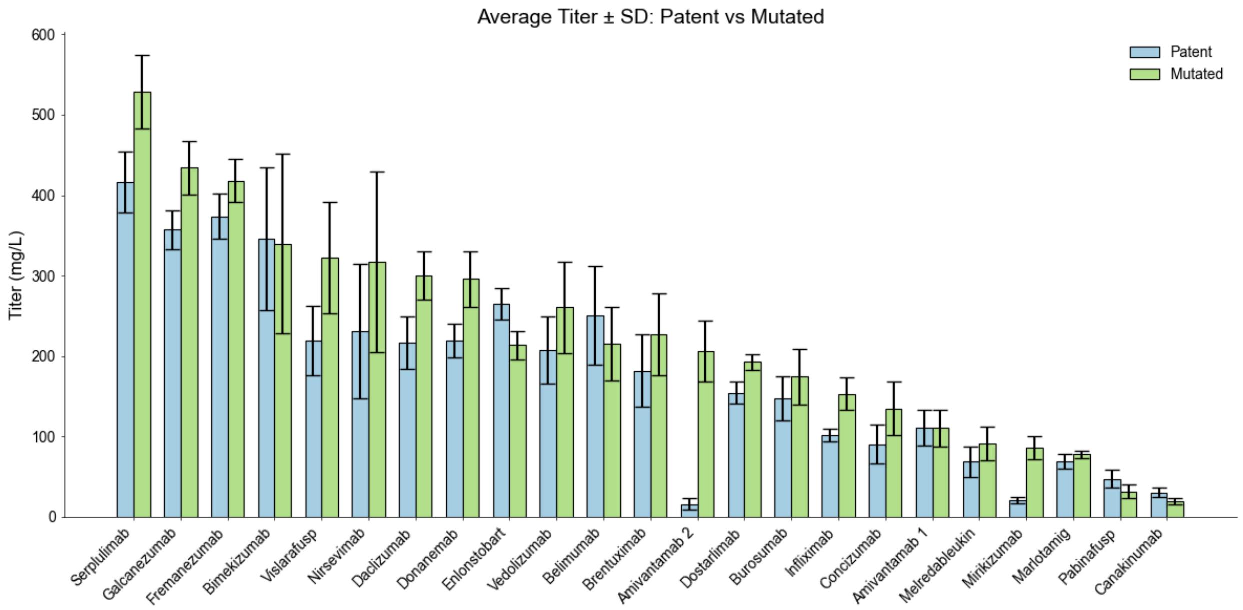


**Supplementary Figure 4: Raw expression titer values across the panel of 23 biotherapeutics.**

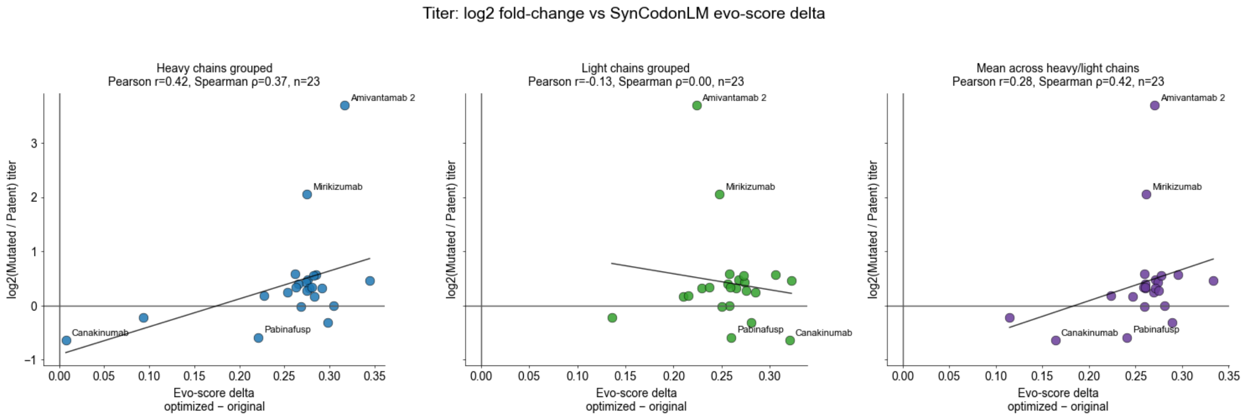


**Supplementary Figure 5: Evo score change from patented sequences per chain type and its effect on expression titer.** A stronger positive correlation is visible in the left graph than middle – heavy chain evo score deltas versus titer improvement, likely due to heavy chain being the limiting factor in transient transfections where two light chain plasmids are added for every heavy chain plasmid, ensuring proper folding of dimerized antibodies. Well optimized light chain plasmids may take translational machinery away from the intrinsically limited – heavy chain.

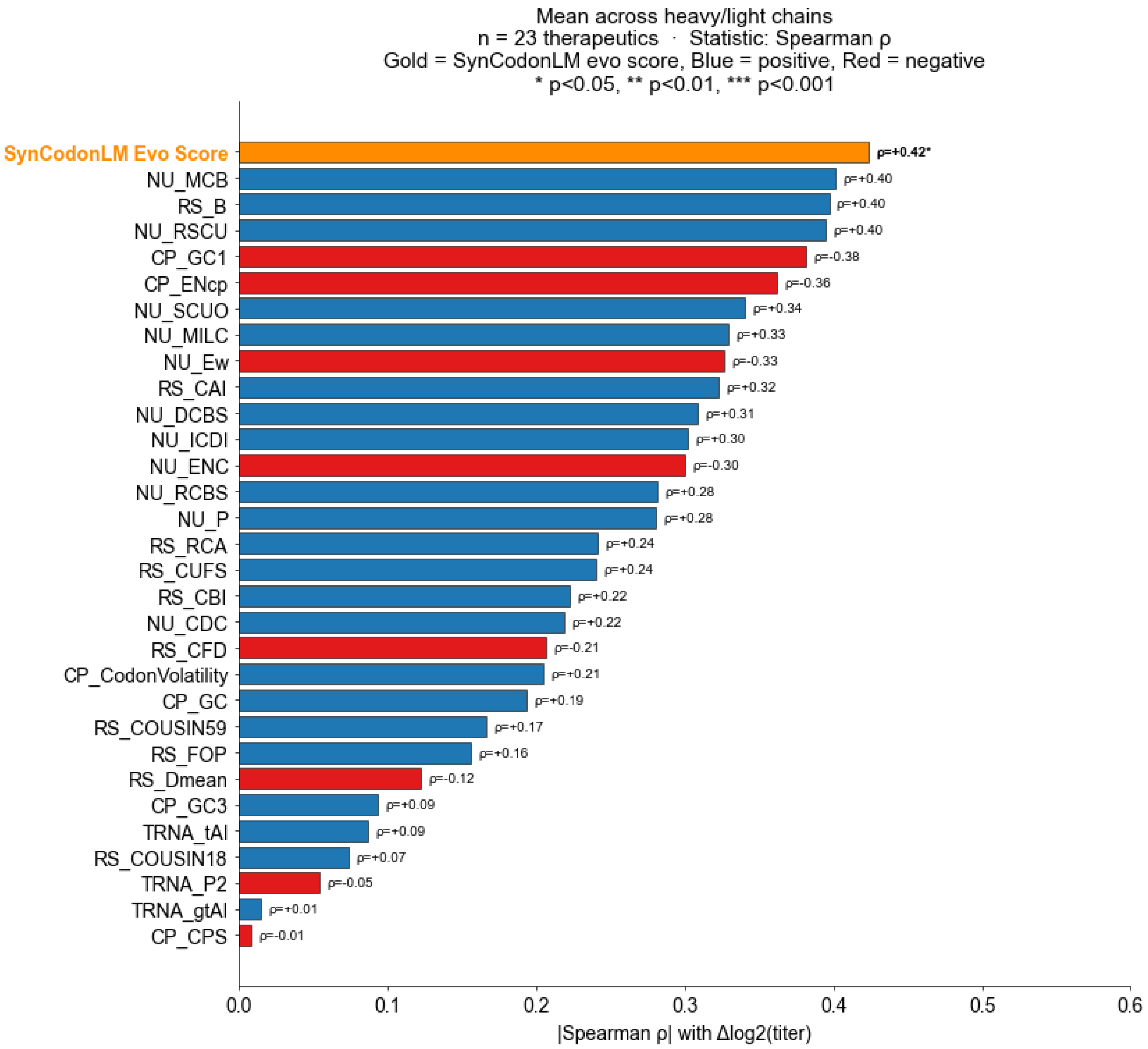


**Supplementary Figure 6: All heuristic statistical descriptors from Figure 5a., and their correlation with Δ$\log_2$(titer).** SynCodonLM Evo score is the only statistically significant descriptor of translational efficiency change.

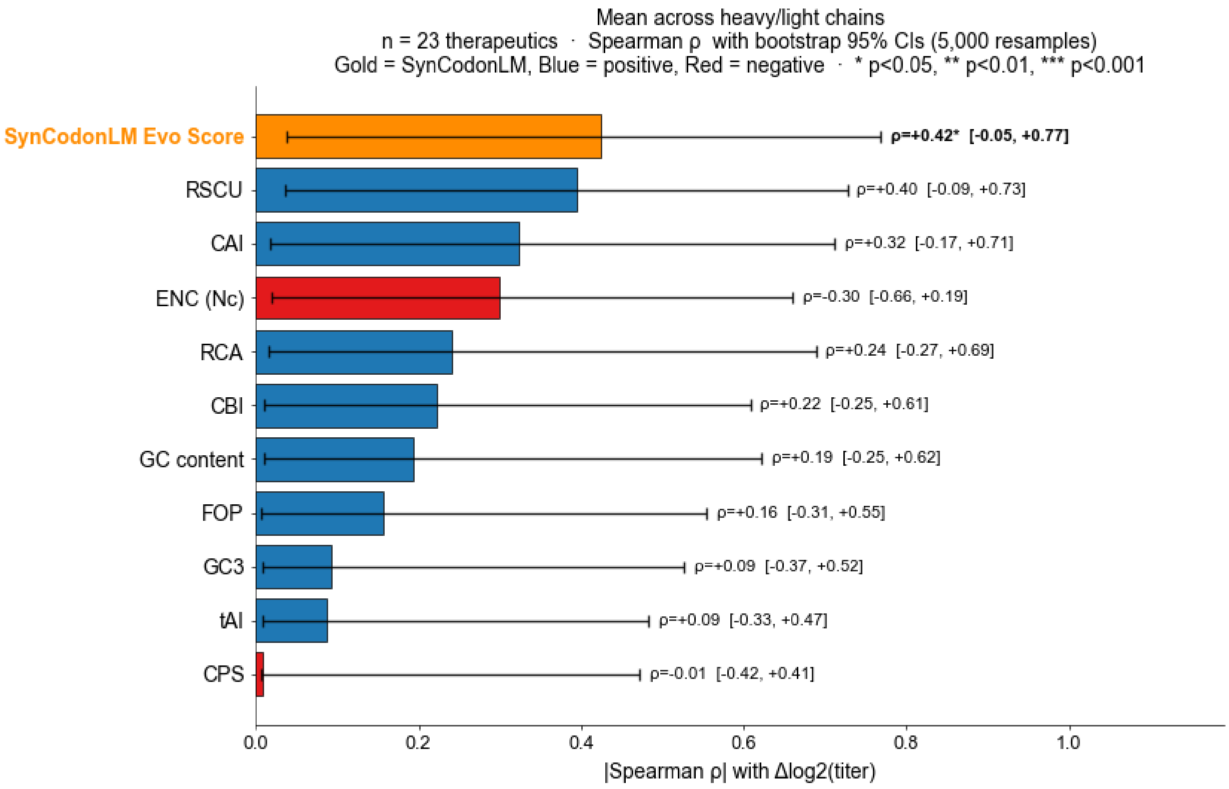


**Supplementary Figure 7: Bootstrap confidence intervals for correlations between descriptor changes and titer changes.** To assess the robustness of the correlation between per-metric Δ and $\Delta\log_2$(titer), we performed nonparametric case-resampling bootstrap analysis. For each metric, paired (Δmetric, $\Delta\log_2$(titer)) observations across the 23 therapeutics were resampled with replacement 5,000 times, and Spearman's rank correlation coefficient (ρ) was recomputed on each resampled dataset. Ninety-five percent confidence intervals were derived as the 2.5th and 97.5th percentiles of the resulting bootstrap distribution (percentile method).

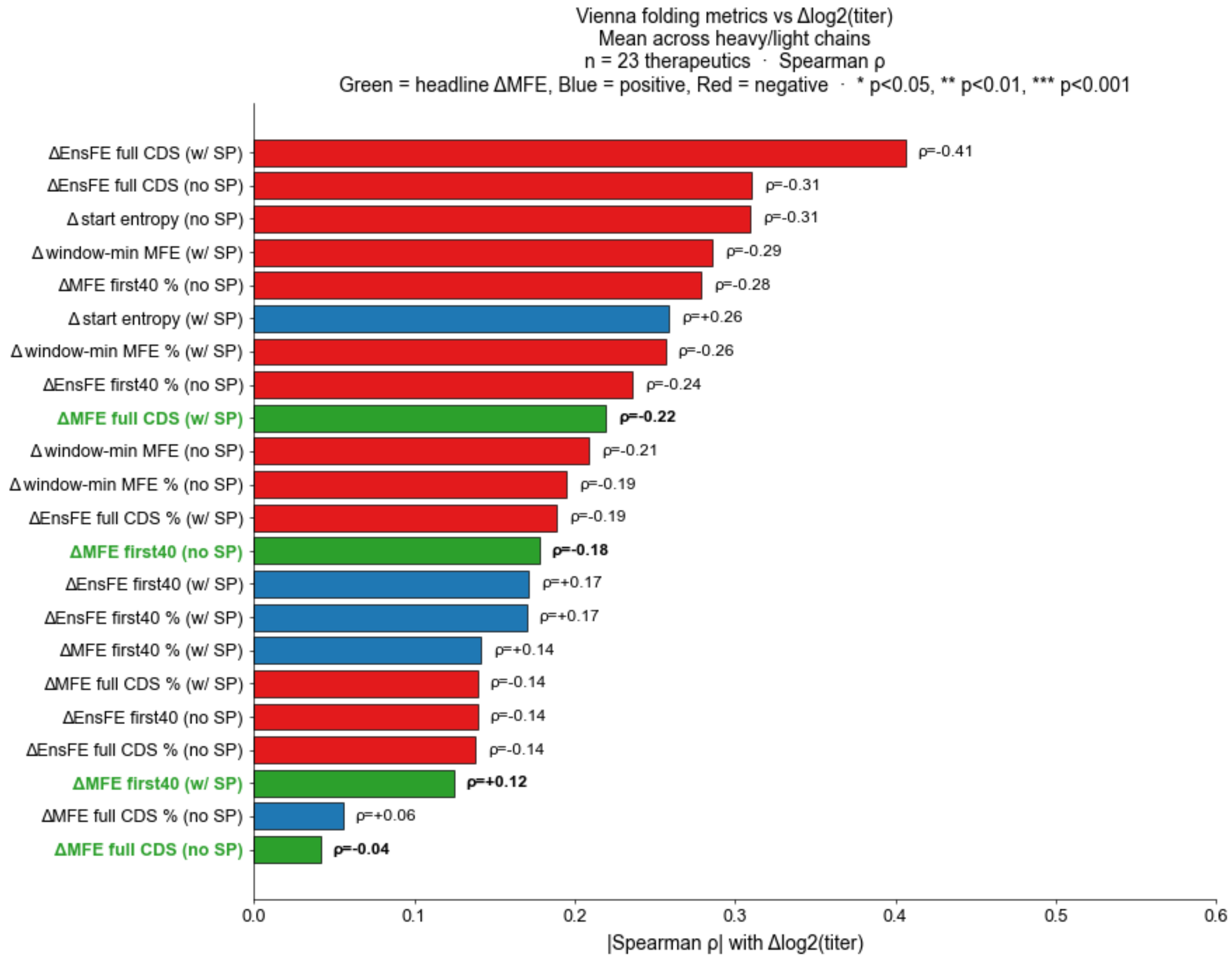


**Supplementary Figure 8: ViennaRNA-derived mRNA-structure descriptors show correlations with $\Delta\log_2$(titer) comparable to, but generally weaker than, the SynCodonLM Evo score.**

Mean absolute Spearman correlation (|ρ|, averaged across heavy- and light-chain-level computations) between the paired change in each ViennaRNA-derived mRNA-structure descriptor (Δ = refined − parental) and $\Delta\log_2$(titer) across 23 clinical-stage antibody-based therapeutics; signed ρ values are annotated on each bar. Descriptors are shown for both folding-input configurations: with the fixed signal peptide and stop codon included (w/SP) and with the mature coding sequence only (no SP). Green highlights the four headline ΔMFE descriptors commonly reported in the codon-optimization literature; blue and red denote positive and negative correlations for the remaining descriptors, respectively. Inclusion of the signal peptide modestly altered descriptor rankings and strengthened correlations for several descriptors, particularly ensemble free-energy metrics.

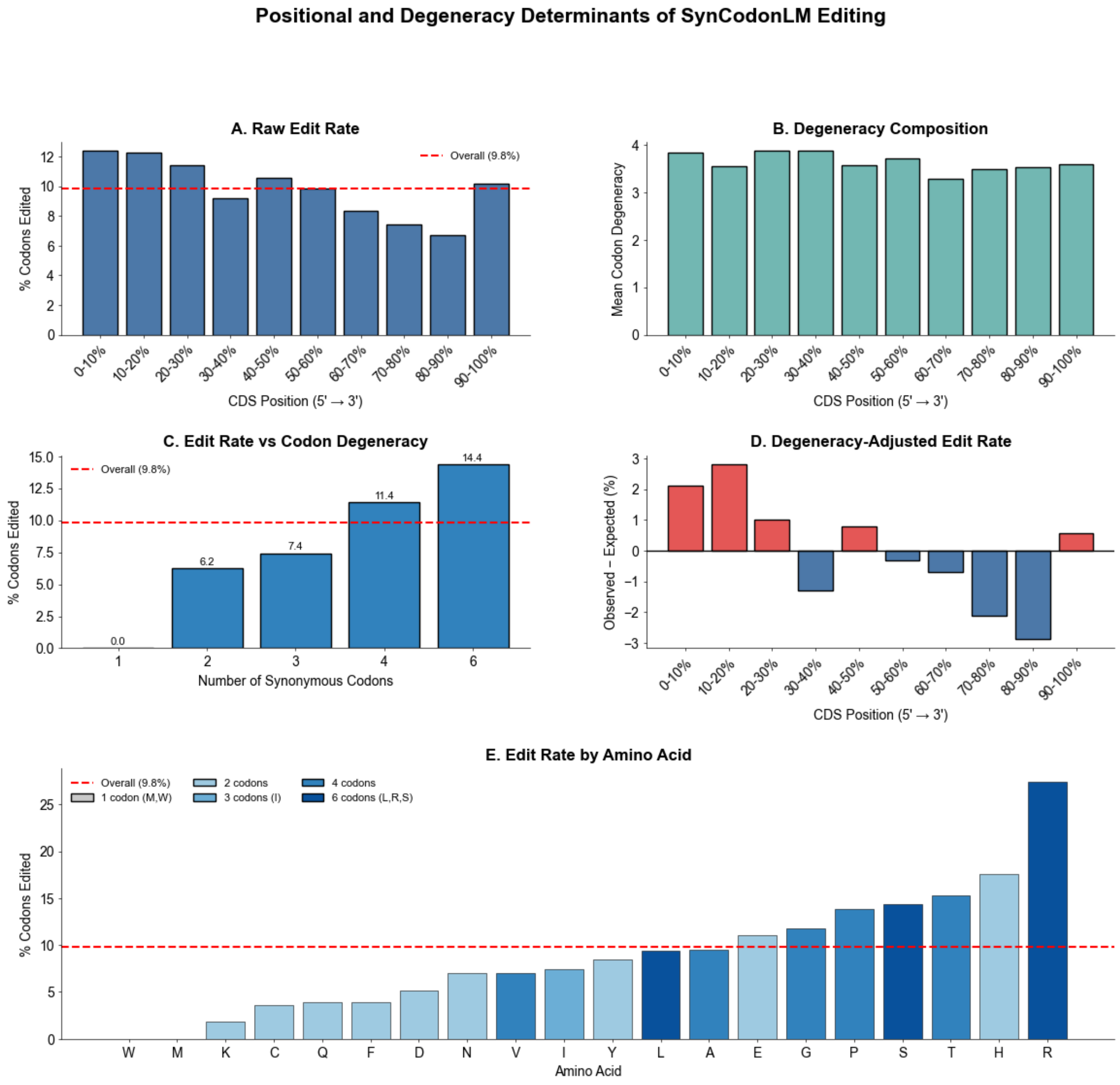


**Supplementary Figure 9: Positional and degeneracy determinants of SynCodonLM editing.** Synonymous codon edits proposed by SynCodonLM across the antibody coding-sequence panel are decomposed by CDS position, codon degeneracy, and amino acid identity. In all panels the red dashed line marks the overall edit rate (9.8%). (A) Raw edit rate (percent of codons edited) binned into deciles along the coding sequence from 5′ to 3′, showing elevated editing in the 5′ region and a decline toward the 3′ end. (B) Mean codon degeneracy (number of synonymous codons per residue) within each positional bin, confirming that codon-family composition is approximately uniform across the CDS and therefore does not by itself explain the positional trend in (A). (C) Edit rate stratified by codon degeneracy class (1, 2, 3, 4, or 6 synonymous codons). Editing increases monotonically with degeneracy, from 0% for single-codon residues (Met, Trp) to 14.4% for six-fold degenerate residues, indicating that the model preferentially edits positions where more synonymous choices are available. (D) Degeneracy-adjusted edit rate, expressed as the observed minus expected editing percentage after accounting for the local codon-degeneracy composition in each bin. Positive values (red) indicate editing above the degeneracy-based expectation and negative values (blue) below it; the residual 5′ enrichment persists after adjustment, showing that positional bias is not solely a consequence of codon degeneracy. (E) Edit rate by amino acid, ordered by increasing editing frequency and colored by codon degeneracy class (see legend). Highly degenerate residues (Arg, Ser, Leu, His) are edited most frequently, while single-codon residues (Met, Trp) are never edited, consistent with the degeneracy dependence shown in (C).

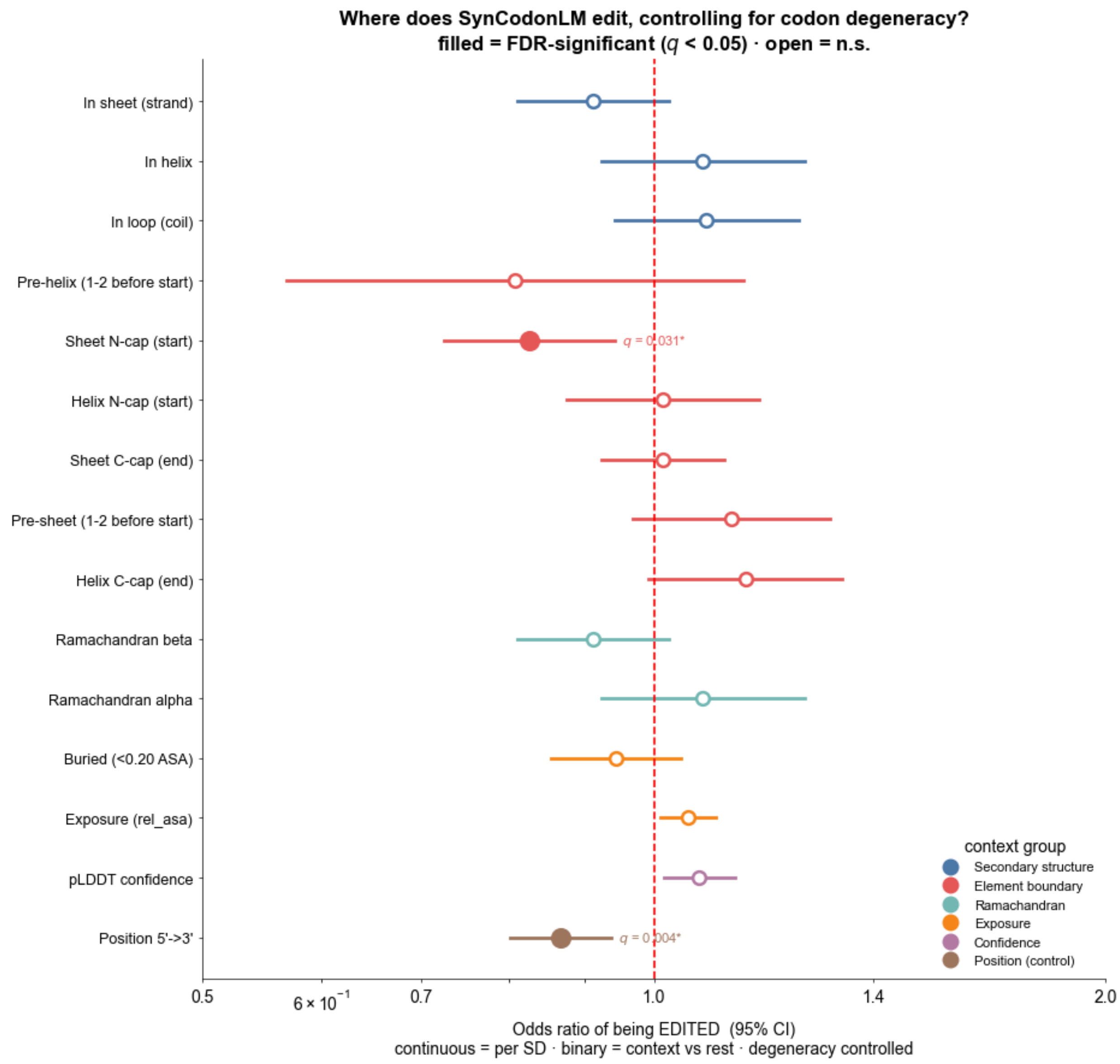


**Supplementary Figure 10: Exploring the structural context of SynCodonLM editing after controlling for codon degeneracy.**

Various structural or positional contexts were each tested in a separate logistic regression predicting whether a codon was edited, with codon degeneracy included as a standardized covariate and standard errors clustered by antibody chain (edited ~ context feature + degeneracy, cluster-robust by chain). Points show the odds ratio (OR) of a codon being edited in each context and horizontal lines the 95% confidence interval; binary features are expressed as context versus all other residues, and continuous features (relative solvent accessibility, pLDDT, and normalized 5′→3′ position) as the OR per standard deviation. The vertical red dashed line marks OR = 1 (no association): OR > 1 suggests editing may be enriched in that context and OR < 1 that editing may be depleted. Filled markers denote associations that remain significant after Benjamini-Hochberg FDR correction across the full panel ($q < 0.05$); open markers are not significant. Contexts are grouped by category (secondary structure, element boundary, Ramachandran basin, solvent exposure, model confidence, and sequence position; see legend) and ordered within group by odds ratio. After degeneracy control, two features appear to reach significance following FDR correction: codons at the N-terminal start (N-cap) of β-sheet strands may be edited somewhat less often than expected (OR < 1, $q = 0.031$), and editing appears to decline from the 5′ to the 3′ end of the coding sequence (OR < 1 per SD of position, $q = 0.004$). Most secondary-structure, Ramachandran, exposure, and confidence contexts show odds ratios close to 1 and are not significant, which may suggest that SynCodonLM editing in this panel is largely associated with codon degeneracy and 5′→3′ position rather than with local structural context, though a possible reduction in editing at strand starts warrants further investigation.
